# Undoing of firing rate adaptation enables invariant population codes

**DOI:** 10.1101/2024.09.26.614457

**Authors:** S.C. Brandão, L. Ramirez, P. Züfle, A.M. Walter, M. Silies, C. Martelli

## Abstract

Neural adaptation supports coding efficiency by tuning responses to prevailing stimulus statistics. However, when information is represented by neural populations, adaptation of individual units could degrade behaviorally relevant signals. Here we investigate how the fly olfactory system implements adaptation in Olfactory Receptor Neurons (ORNs) and the consequences for combinatorial coding in downstream circuits. We show that adaptation of ORN firing rate is compensated at the axon terminal, where calcium transients remain background-invariant through inhibitory presynaptic feedback. Background invariance requires an adaptation strategy that shifts ORN response amplitude rather than sensitivity, diverging from efficient coding principles in single neurons. This property supports contrast encoding in ORN populations necessary for background compensation across the glomeruli. Downstream, the modulation of presynaptic Unc13 proteins maintains postsynaptic projection neurons responses to ON stimuli background invariant. We identify a new coding strategy where olfactory neuronal populations encode asymmetrically contrast information by implementing circuit computations that compensate peripheral firing rate adaptation.

## Introduction

Neural adaptation is an essential strategy of sensory systems that allows the encoding of stimuli over a range of intensities exceeding the limits of the individual sensory cell^1^. To do so, neurons tune their stimulus-response function to the relevant statistics of the stimulus^2^. For instance, efficient coding theories at the single-neuron level predict that a shift in neuron sensitivity that matches the mean stimulus optimizes the encoding of relevant information within the neuron’s dynamic range^3^. Such shifts in sensitivity have been observed in several sensory systems, from audition to vision, in artificial and natural conditions^4–6^. However, much less is known about the role of adaptation in odor coding. The olfactory system is by design endowed with a large array of sensory receptors with different affinities to the same odorant^7,8^. When the stimulus intensity (concentration) increases, the more sensitive receptors saturate while new ones with lower affinity are recruited. Therefore, the system as a whole is rarely out of its broad dynamic range. While the benefits of adaptation in single neurons have been widely discussed^9,10^, the benefits of single neuron adaptation for population coding require a deeper understanding. Studies in the visual cortex suggest that neuronal adaptation can improve coding accuracy at the population level in a stimulus-dependent manner^11^. Here we investigate the function and potential benefits of adaptation in populations of olfactory neurons.

In both vertebrates and invertebrates, the firing rate of Olfactory Receptor Neurons (ORNs) decreases upon persistent stimulation^12–18^, inducing a change in their response curve. Yet, the coding function of this adaptive change remains unclear. On one hand, a decrease in ORN response could lead to a decrease in perception of the odor stimulus. However, the relationship between ORN adaptation and decreased stimulus perception is so far correlative^19^ since a decreased behavioural response to persistent odour stimuli could derive from plasticity (e.g. habituation^20–22^) in higher brain areas rather than from decreased receptor sensitivity. On the other hand, adaptation of the sensory neurons could play a role in odor-driven navigation. Behavioral studies in *Drosophila* suggest that flies can implement a chemotactic strategy for odour-guided searching behaviour^23,24^. In single cell organisms, such as bacteria, chemotaxis relies on receptor adaptation for contrast computations^25^. Such contrast computations could in principle also occur in the ORNs^26,27^, although the contrast sensitivity of these neurons across a physiological range of concentrations is marginal^28,29^. Importantly, contrast computation in single ORNs prior to the encoding of odor identity across the ORN population could interfere with information transmission in downstream circuits.

To understand the role of peripheral ORN adaptation it is necessary to analyze odor coding principles in downstream pathways. ORNs expressing the same chemosensory receptor send axons to the same olfactory glomerulus, where they connect to projection neurons (PNs) in insects and mitral/tuft cells (M/T cells) in vertebrates^30^. Synaptic depression in these first order connections^31^ and lateral inhibition^32^ mediated by local neurons (LNs) modulate odor representations in the insect antennal lobe^33^ (AL) and in the vertebrate olfactory bulb^34^. However, a role of these two circuit mechanisms in adaptation has not been investigated so far.

Here we combine computational and experimental approaches in *Drosophila* to understand how ORN adaptive properties shape odor representations in populations of second-order olfactory neurons, the PNs. We start with a computational analysis to show that the odor information encoded in populations of ORNs depends on adaptation properties of single neurons. Specifically, an adaptive change in response amplitude, rather than in response sensitivity, supports the encoding of both contrast and intensity in a population of ORNs. We show that adaptation by a change in response amplitude is required to drive the proper amount of feedback inhibition in the ORN presynapses to maintain calcium responses invariant to background stimuli, undoing the effect of firing rate adaptation. This invariance is specific for ON stimuli (increase in concentrations compared to the background) creating an asymmetry between ON and OFF odor representations. Such asymmetric invariance persists in postsynaptic PNs and is disrupted by mutation of a modulatory domain of Unc13, suggesting a role of presynaptic plasticity in achieving the response invariance. We propose that adaptation in ORN firing rates enhances the separation of ON and OFF stimuli in the downstream population of PNs, while allowing a background-invariant representation of odor identity for ON stimuli which requires the encoding of contrast information in ORN populations.

## Results

### Single neuron adaptation shapes odor information coding in ORN populations

Adaptation has often been described in both peripheral and central sensory neurons as a mechanism that tunes neuron sensitivity to the stimulus statistics. For example, adaptation to an increasing background luminance leads to a shift in the response curve of fly photoreceptors towards higher luminances^35^. This shift is consistent with the efficient coding hypothesis^10^ and implies that the neuron encodes contrast rather than absolute luminance. ORNs in both insects and vertebrates do not seem to follow the same adaptive strategy as visual neurons since they do not exhibit a shift in sensitivity (in the stimulus domain), but rather a decreased response (in the activity domain) after adaptation to an odor background (reviewed in ^29^). An important difference between vision and olfaction is that stimulus sensitivity across ORNs is highly heterogeneous due to the expression of large odorant receptor repertoires, leading to an early combinatorial encoding of odor information by the ORN population^7,8^ (**Fig. 1a**). To understand how single neuron adaptation shapes odor coding at the population level, we simulated the response of a population of ORNs under three different strategies for background adaptation (**Fig. 1b**): I) sensitivity-shift (core mechanism in contrast-sensitive visual neurons), II) amplitude-compression and III) amplitude-shift, the latter two being plausible based on measurements in ORNs^14,29^. For each scenario, we simulated the population response to increasing concentrations of a single odor presented in isolation or after adaptation to a background of the same odor. For all three cases, we described the population responses in a low-dimensional manifold focusing on its capability to encode either stimulus intensity or contrast (relative to the background). If single ORNs adapt by shifting their sensitivity (**Fig. 1c**), the population responses lie on a one-dimensional manifold where the stimulus intensity strongly correlates with the first principal component, which explains most of the response variance across all backgrounds (**Fig. 1d**). Background adaptation leads to similar representations of iso-intensity stimuli (**Fig. 1d**, black dots). Conversely, the encoding of contrast information in this population is ambiguous, as the same contrast can be represented by very different representations (**Fig. 1e**, black dots). Therefore, an adaptive strategy that allows the encoding of contrast in single neurons leads to population responses that retain mostly information about intensity. On the contrary, in a population of neurons that adapt their response amplitude by a compression or a shift, odor representations lay on a 2-dimensional manifold (**Fig. 1f, i**), allowing the parallel encoding of both intensity and contrast information (**Fig. 1g-h and j-k**, black dots). These results generalize across different update rules for the parameters (see Methods).

**Figure 1.**
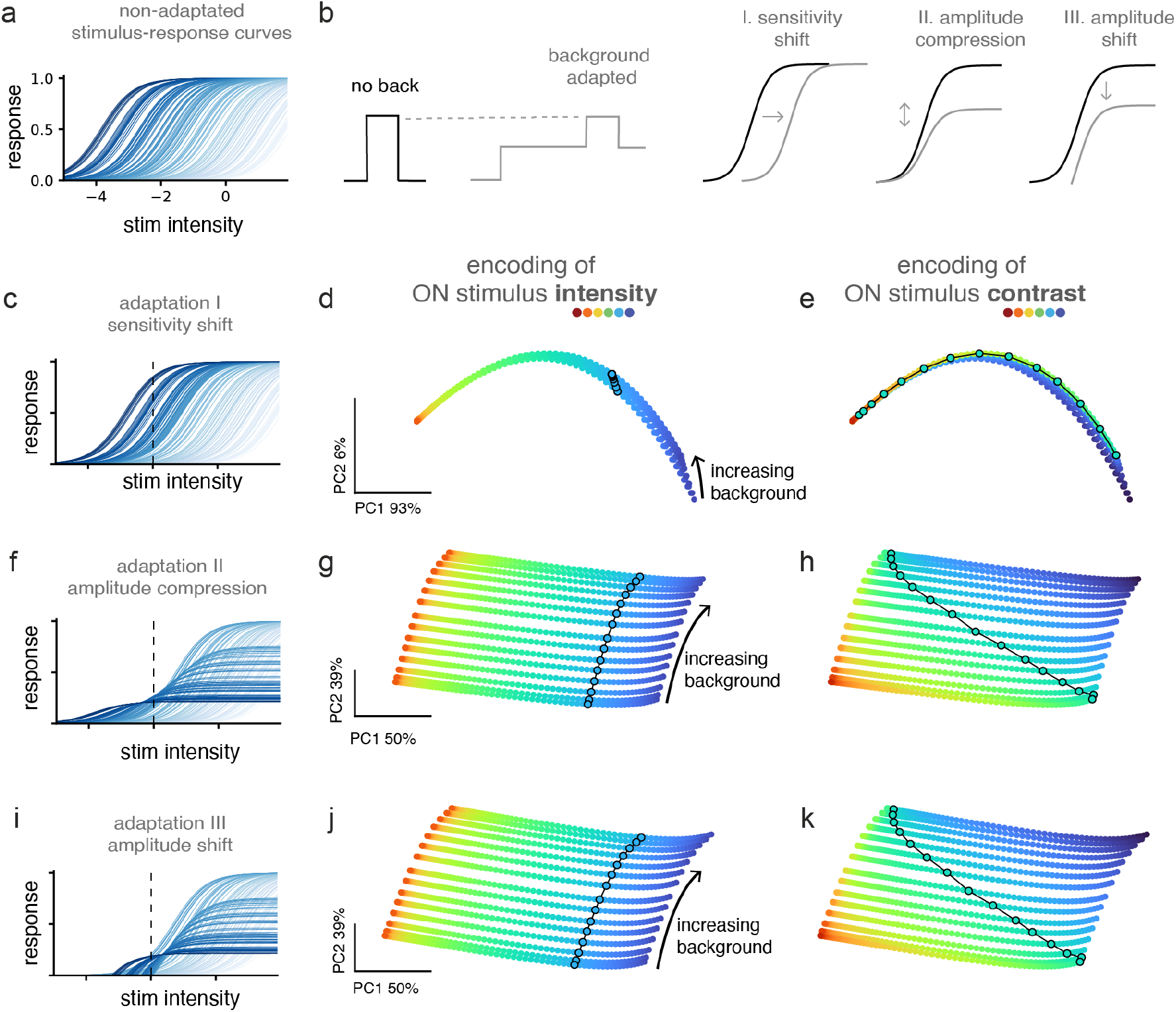
Encoding of intensity and contrast in neuron populations with different adaptive properties. **a)** Simulated stimulus response curves for a population of ORNs with different stimulus sensitivities (stimulus in log scale). **b)** Three types of background-induced adaptation for individual neurons, implemented as a change in a specific parameter for each adaptation case (see Methods). **c)** Response curves of the same neurons as in a) after adaptation to a background stimulus indicated by the dotted line. Adaptation to the background induces a shift of the response sensitivity towards higher stimulus intensities. **d)** 2D-manifold of the population responses to stimuli of different intensity (color coded) in different background adapted conditions. Black circles highlight an iso-intensity response curve across all backgrounds. **e)** Same data as in **d)** but color coded by stimulus contrast. Black circles highlight an iso-contrast response curve across all backgrounds. **f-g-h)** as in **c-d-e)** for a population of neurons that adapt by a compression in amplitude. **i-j-k)** as in **c-d-e)** for a population of neurons that adapt by shifting their response amplitude.

Altogether, we find that adaptation strategies at the single neuron level determine how odor information is represented by the population. Importantly, a strategy that is commonly used in other sensory systems (strategy I) performs poorly at encoding stimulus features such as contrast at the population level when compared to adaptive strategies that modify response amplitude (strategies II and III). This counter-intuitive result has a simple explanation. When adaptation induces a shift in sensitivity, the total amount of activity generated by the ORN population is the same across background conditions. Instead, when adaptation leads to a change in response amplitude (a compression or shift), the total activity in neural space decreases with background intensity. This shifts the representational space, creating a dimension to encode contrast as combination of background and stimulus intensity. In the following we ask whether and how this coding scheme is passed to or used by downstream neurons.

### ORNs transform adapted firing rate responses to ON odor stimuli into background-invariant calcium representations

To understand the computational role of ORN firing rate adaptation, it is essential to examine how these signals are processed in the olfactory glomeruli, where ORNs project their axons (**Fig. 2a**). We have previously characterized the response dynamics of ORNs^14,36^, which we briefly summarize here. ORNs exhibit phasic firing rate responses to an increase in odor concentrations, with activity peaking shortly after stimulus onset and subsequently relaxing to an adapted steady-state. In the adapted state the neuron’s response to subsequent odor stimuli is weaker (**Fig. 2b**, data from^36^). This form of background-dependent adaptation, typical of sensory neurons, is observed at the dendritic terminals within the antenna (**Fig. 2a, b**). The dendritic firing activity drives calcium transients in the axon terminals of ORNs within specific glomeruli of the antennal lobe (**Fig. 2a**). We have previously shown that these presynaptic calcium signals exhibit sustained, rather than phasic, dynamics (**Fig. 2c**) and, importantly, that their amplitude remains invariant to background odor levels despite adaptation of the firing rate (**Fig. 2c** and more data in^36^). Therefore, when odor stimuli are presented in isolation, presynaptic calcium transients, assayed by fluorescence indicators, depend linearly on firing rate, but this relationship is altered by background adaptation (**Fig. 2d**, data adapted from^36^). These findings led us to hypothesize that presynaptic calcium dynamics and therefore synaptic release are regulated in an activity-dependent manner.

**Figure 2.**
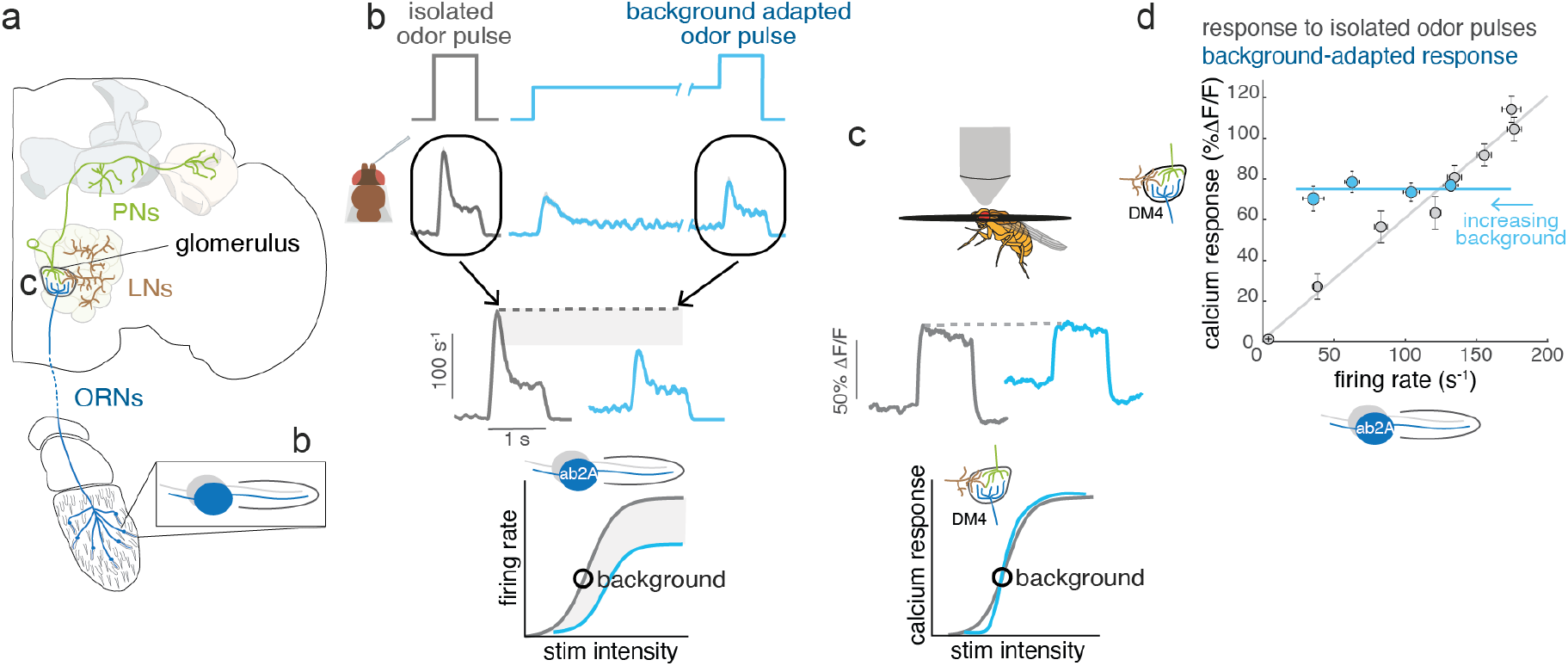
The undoing of firing rate adaptation at central synapses. **a)** Schematics of the olfactory pathway. Olfactory Receptor Neurons (ORN) located in the antenna extend their axons to a specific glomerulus in the antennal lobe and make synapses onto Local Neurons (LNs) and uniglomerular Projection Neurons (PNs). PNs’ axons innervate the Lateral Horn and the calix of the Mushroom Body. The labels (b and c) indicate the location of recordings of the corresponding panels. **b)** *Top:* Schematics of the stimulus protocol and firing rate response of the ab2A ORNs in control and background adapted conditions (data adapted from^36^). *Bottom:* schematics of the ORNs firing response in control and background-adapted conditions. The circle indicates the intensity of the background stimulus. **c)** *Top:* ORN calcium responses induced by an odor puff in control and adapted conditions measures from the ab2A ORN axon terminals in the corresponding glomerulus DM4. *Bottom:* schematics of the ORN calcium response curve showing background-invariant responses. **d)** Mean firing rate and mean calcium responses for odor puffs of different intensity tested in adapted (blue) and non-adapted (gray) conditions (data from^36^). The lines are linear regressions. Error bars indicate SE.

### Activation of a single glomerulus is sufficient for background-invariant ORN presynaptic activity

Odor stimuli can recruit more than one ORN type, raising the question of whether the background invariance of ORN presynaptic responses depends on multi-glomerular activation. We used an optogenetic approach to restrict activity to a single ORN type (expressing the odorant receptor Or42b) targeting the glomerulus DM1 (**Fig. 3a**). Using a combination of background and test stimuli, we show that, as with odors, ON stimuli elicited the same response irrespective of background, while OFF stimuli always lead to drops in calcium below the non-adapted response (**Fig. 3b, c**). We refer to this encoding property of the olfactory system as “asymmetric background invariance*”*. We further measured firing rates from the ab1 sensillum that contains the Or42b-ORNs to verify that the firing rate responses are actually adapted to the stimulus background (**Fig. 3d**). Optogenetic activation leads to lower firing rates and a smaller dynamic range compared to odor stimuli (response saturates at around 40 Hz rather than the usual 250 Hz). Yet, as with odors, background adaptation leads to a striking change of the dose-response curve for both ON and OFF stimuli (**Fig. 3d**). We conclude that presynaptic calcium is regulated such that both uni-glomerular (**Fig. 3**) and multi-glomerular (**Fig. 2**) odor representations are normalized to achieve asymmetric background invariance.

**Figure 3.**
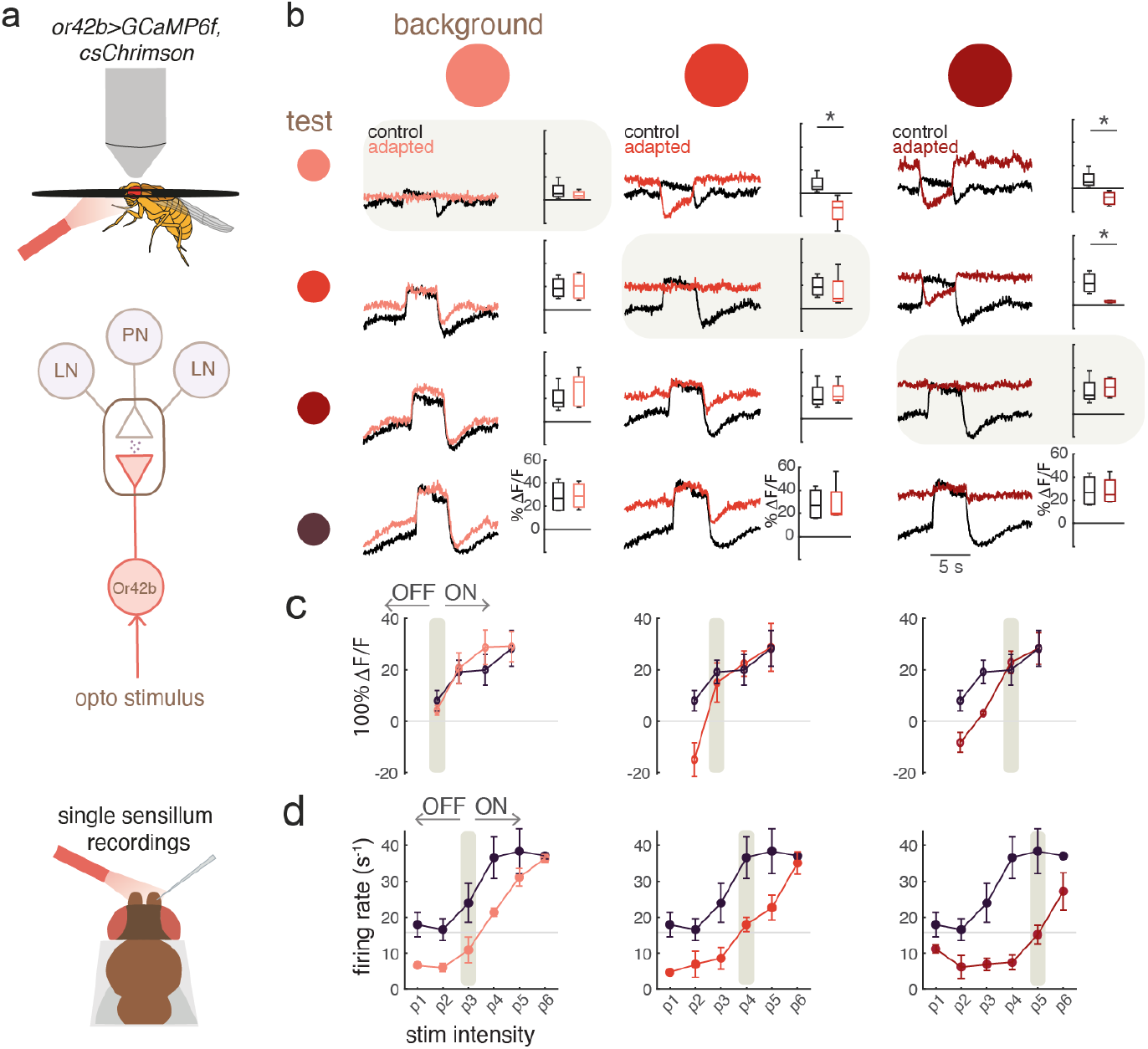
Asymmetric background-invariance in ORN upon optogenetic activation. **a)** Schematics of the experimental protocol. **b)** Mean calcium responses from ORN axon terminals in the glomerulus DM1 in control (black, no background) and adapted (shades of red) conditions. Columns represent different background intensities, rows different intensities of the test stimulus. Test intensities are: 0.01, 0.03, 0.055, 0.08 μW/mm^2^ (from light to dark red). The plots on the diagonal (gray quadrants), corresponds to the same intensity of the test and background stimuli. Quadrants above this diagonal, are OFF stimuli and quadrants below are ON stimuli. Boxplots show median, quartiles and min/max values of the response averaged in 3 sec after stimulus onset (N=4-5, *p<0.05). **c)** Mean response plotted as a function of the test stimulus intensity; error bars indicate SE. **d)** Same as **c)** but for firing rate responses (N=3). A similar range of stimuli was used for the electrophysiology (p1-p6 = 0.06, 0.1, 0.2, 0.4, 0.7, 0.92 μW/mm^2^). Gray boxes indicate the intensity of the background stimulus.

### Homeostatic feedback regulates ORN presynaptic calcium responses

Next, we investigated the potential mechanisms that compensate the firing rate adaptation and achieve ON background invariance. Specifically, we asked whether ORN presynaptic calcium is regulated in a cell-intrinsic manner or through a circuit computation. To distinguish between these two cases, we expressed the temperature-sensitive dynamin mutant shibire^ts^ in Or42b-ORNs to block neurotransmitter release and measured odor responses at the same presynapses (**Fig. 4a**). To capture slower adaptive processes, we presented the test odor 8 consecutive times (**Fig. 4b**). As expected, flies of the control genotype showed tonic and background-invariant responses (**Fig. 4c**). In contrast, blocking synaptic release led to more transient and background dependent responses (**Fig. 4d-f**), reminiscent of firing rate adaptation (**Fig. 2a, b**). Similar results were obtained for the glomerulus DL1 (**Fig. S41a-d**). Therefore, ORN presynaptic calcium depends on other circuit components to compensate the change in firing rate induced by background adaptation. Importantly, lateral inputs from other glomeruli, activated by this odor but unaffected by the manipulation (e.g. from DM4), are not sufficient to compensate adaptation of the DM1-ORNs. Together with the optogenetic manipulations previously discussed, this data provides further evidence for an intra-glomeruli mechanism.

**Figure 4.**
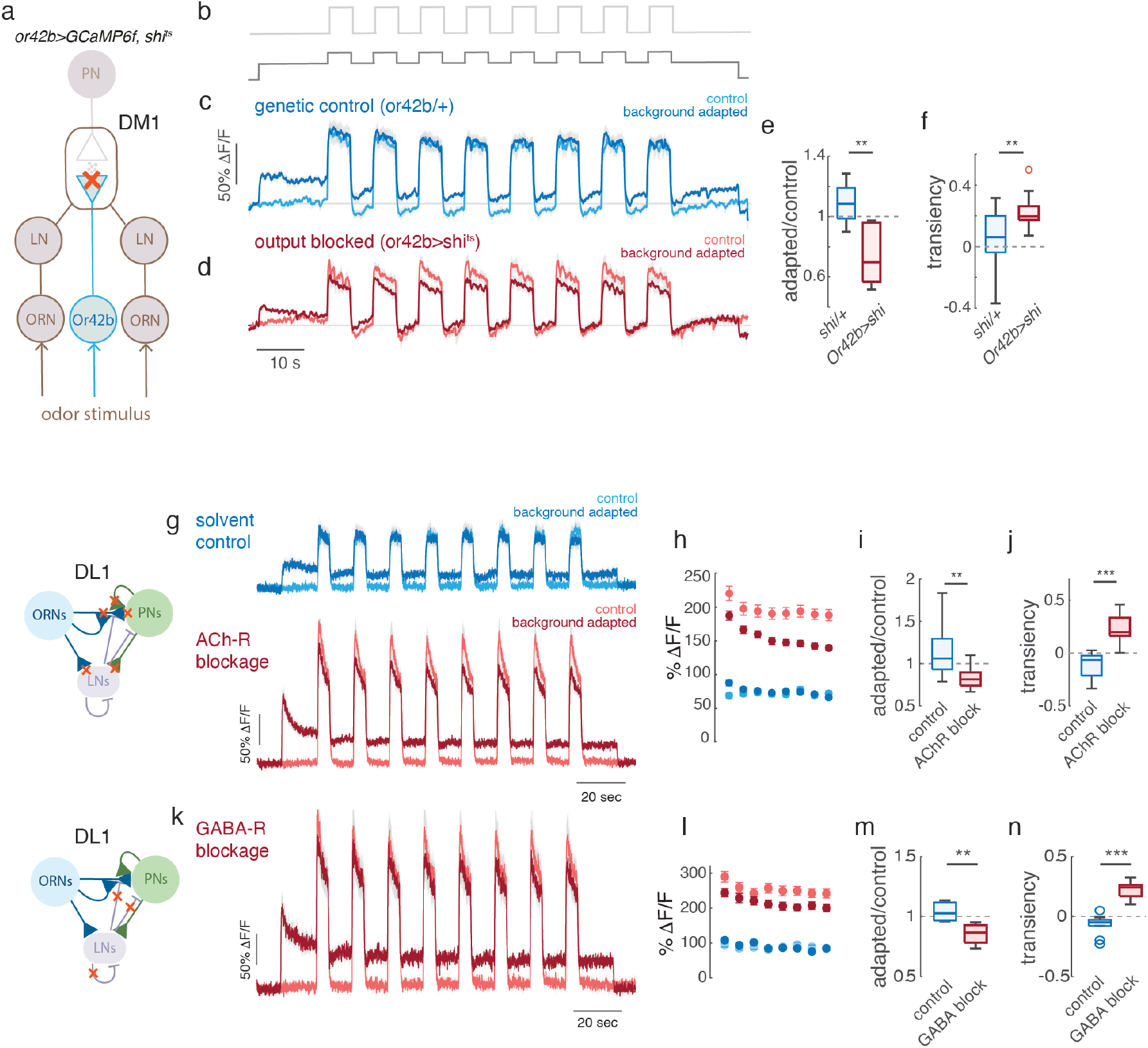
An input gain control mechanism maintains ORN presynaptic calcium background-invariant. **a)** Genetic blockage of neurotransmitter release from Or42b-ORNs. **b)** Stimulation protocol in control (light gray) and background adapted conditions (dark gray). **c)** Calcium response of the DM1 glomerulus in genetic control flies showing background-invariant responses and **d)** calcium response in flies with blocked ORN synapses showing background-dependent responses to methyl acetate (liquid dilution of 10^−4^, background airflow dilution C5 and pulse C6, see **Table S3**). **e)** Relative adaptation measure as the response ratio in adapted and control conditions. **f)** Response transiency measured as the relative change in response at stimulus onset and offset (transiency=1-f_offset_/f_onset_) N=7, 8. **g)** Calcium responses from the DL1 glomerulus before and after drug application. Pure 2-butanone was used in airflow dilutions of C7 for the background and C8 for the pulse. For the blockage of AChR 100 μM atropine + 20 μM tubocurarine + 200 nM imidacloprid was used, N=10. **h)** Mean response to the single pulses, error bars indicate SE. **i-j)** As in **e-f). k-n)** as **g-j)** for GABA-R blockage: 25 μM CGP54626 + 5 μM picrotoxin, N=5. Thick lines indicate mean and shaded areas indicate SE. Box plots indicate median, quartiles, min/max values and outliers. **p<0.01, ***p<0.001, Kruskal-Wallis.

ORN synapses release acetylcholine in the AL. Pharmacological block of acetylcholine receptors (AChRs) also led to transient and background-dependent responses (**Fig. 4g-j**) for different combinations of test and background stimuli (DL1, **Fig. S2a-c**), further indicating that the regulation of the presynaptic calcium is not a cell-intrinsic property and requires a synaptic feedback loop. ORNs make most of their synapses with PNs and LNs. Genetically silencing PN output had no effect on presynaptic calcium in ORNs, excluding an output gain control mechanism (**Fig. S41e-f**). LNs form a diverse population of inhibitory neurons mediating different types of gain control in the AL^33,37^, but their role in the context of background adaptation remains unexplored. Pharmacological block of GABA receptors led to similar effects as blocking of AChRs (**Fig. 4m-p, and Fig. S2a**), suggesting that GABAergic LNs play a major role in tuning background adaptation in this glomerulus.

We observed similar or partial effects in other glomeruli (DM3, DM6 in **Fig. S2b**), but not in all (DL5, **Fig. S2**). Some of this variability might be attributed to differences in drug penetration. However, DL5 recordings were obtained from the same preparation, on the same focal plane and with the same stimuli as for DL1, minimizing potential differences in drug access. The background-invariant response of DL5 ORNs is unaffected by the blockage of either AChRs or GABA receptors (**Fig. S2a**,**b**). Additionally, these neurons display an unusual slow increase in baseline calcium across stimulus repetitions, forming a ramp resistant to drug application. This makes us postulate the existence of a cell-intrinsic mechanisms for the regulation of pre-synaptic calcium in DL5.

Overall, our data indicate that, in many glomeruli, inhibitory inputs on the ORN presynapses are necessary to mediate a form of input gain control that counteracts firing rate adaptation, making responses to ON stimuli background invariant. Some glomeruli might implement a redundant or alternative strategy relying on cell-intrinsic homeostatic control of presynaptic calcium.

### An amplitude-shift adaptation mechanism is required to achieve background invariance through inhibitory feedback

We next asked how inhibitory feedback from the LNs can lead to invariant calcium dynamics across different background conditions. To address this question, we implemented a dynamic model of the ORN presynaptic calcium response where ORN firing rates provide the excitatory inputs while LN drive the inhibitory feedback (**Fig. 5a** and Methods). We show that to achieve response invariance across all background conditions, single ORNs must implement an adaptation mechanism of their firing rate that acts as a differential operator in the response domain (**Eq. 7** in Methods). Strikingly, this mechanism corresponds to the amplitude-shift adaptation mechanism that we showed to preserve both intensity and contrast information at the population level (**Fig. 1i-k**). To further understand the role of firing rate adaptation for background invariance of calcium responses, we modeled ORN calcium responses for the three different adaptation mechanisms in **Fig. 1b** (**Eq. 8** in the Methods). The sensitivity-shift mechanism ensures an invariant calcium response in a test-dependent manner, for which information about both the test and background stimuli are required. As a consequence, calcium responses after adaptation to different backgrounds are invariant only for a specified test stimulus (**Fig. 5b**, dashed line). Similarly, the amplitude-compression mechanism leads to a test-dependent response that ensures invariance only for stimuli much higher than the background (**Fig. 5c**). As formally predicted (**Eq. 7** in Methods), only the amplitude-shift adaptation mechanism ensures invariant calcium responses for all backgrounds and test stimuli (**Fig. 5d**). This current-based dynamical model captures both the firing rate and calcium dynamics but does not explicitly include the divisive normalization implemented by LNs^33^. To better understand the interaction between divisive normalization and background adaptation, we also implemented a stationary model of divisive normalization with an inhibitory input provided by adaptive LN neurons (see Supplementary Material). We show that also in this case background invariance is guaranteed only for an adaptation mechanism that is amplitude dependent - in this case a combination of both shift and compression (**Fig. S3a**).

**Figure 5.**
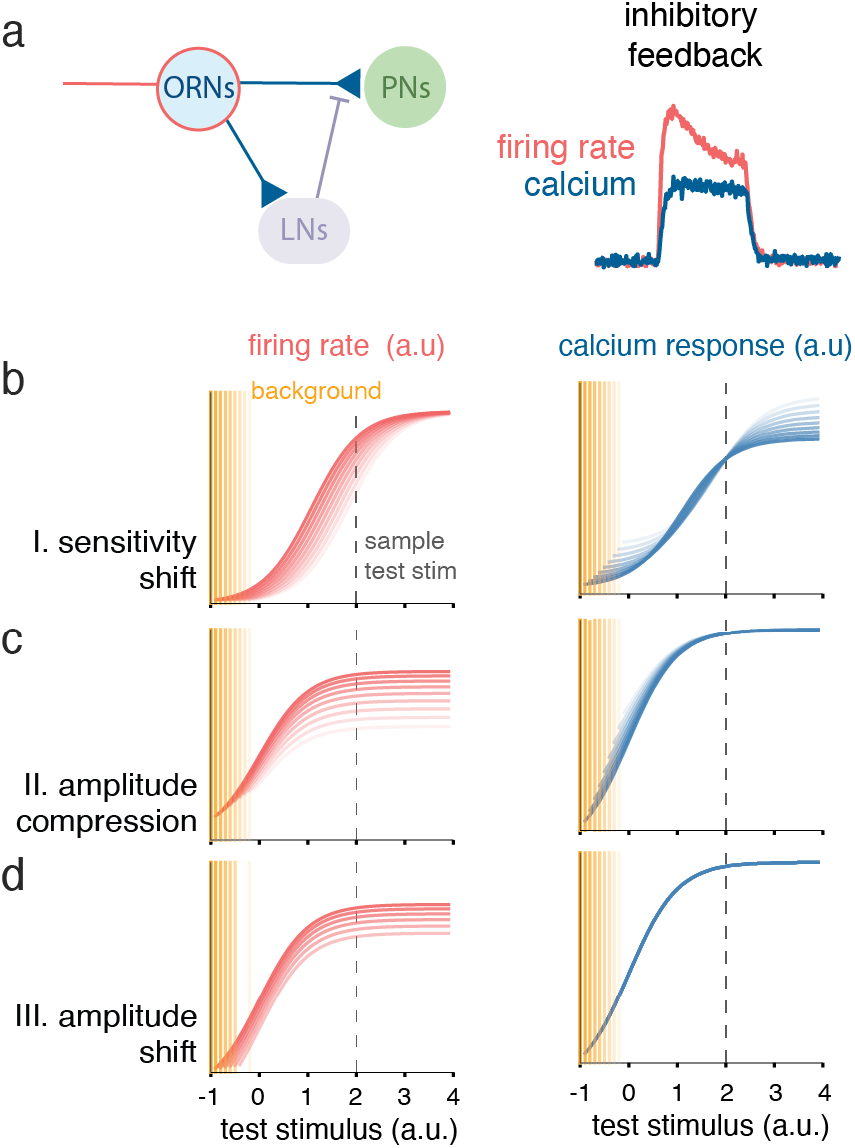
Adaptation mechanisms for background-invariant response. **a)** We consider an inhibitory presynaptic feedback loop with ORN firing rate as input and ORN axonal calcium as output (see Methods and Eqs. (3-8)). **b-c-d)** Simulated firing rate (red) and calcium response (blue) for the three different adaptive functions in Eq. (8): sensitivity-shift, amplitude-compression, amplitude-shift. Different color intensities correspond to different simulated background stimuli indicated by the orange lines. Dotted lines indicate a representative test stimulus.

Altogether, we demonstrated that inhibitory feedback can mediate ON background invariance only if the input firing rate adapts by an amplitude dependent mechanism, highlighting a new computational role of adaptation that deviates from efficient coding predictions at the single-neuron level.

### Olfactory projection neurons inherit a background-invariant representation of ON odor stimuli

To understand the relevance of these axonal computations in ORNs, we investigated whether background adaptation is preserved in the postsynaptic neurons. ORNs make most of their synapses on uniglomerular PNs. ORN-PN synapses undergo depression on both fast^31^ and slow^36^ timescales. We reasoned that even a tonic activation of depressing synapses would result into phasic and adaptive responses in postsynaptic neurons. Indeed, contrary to ORNs, PNs calcium dynamics are phasic (as also shown in ^36^), but surprisingly their responses to odors are independent of background adaptation, as in the ORN axons (**Fig. 6a**).

**Figure 6.**
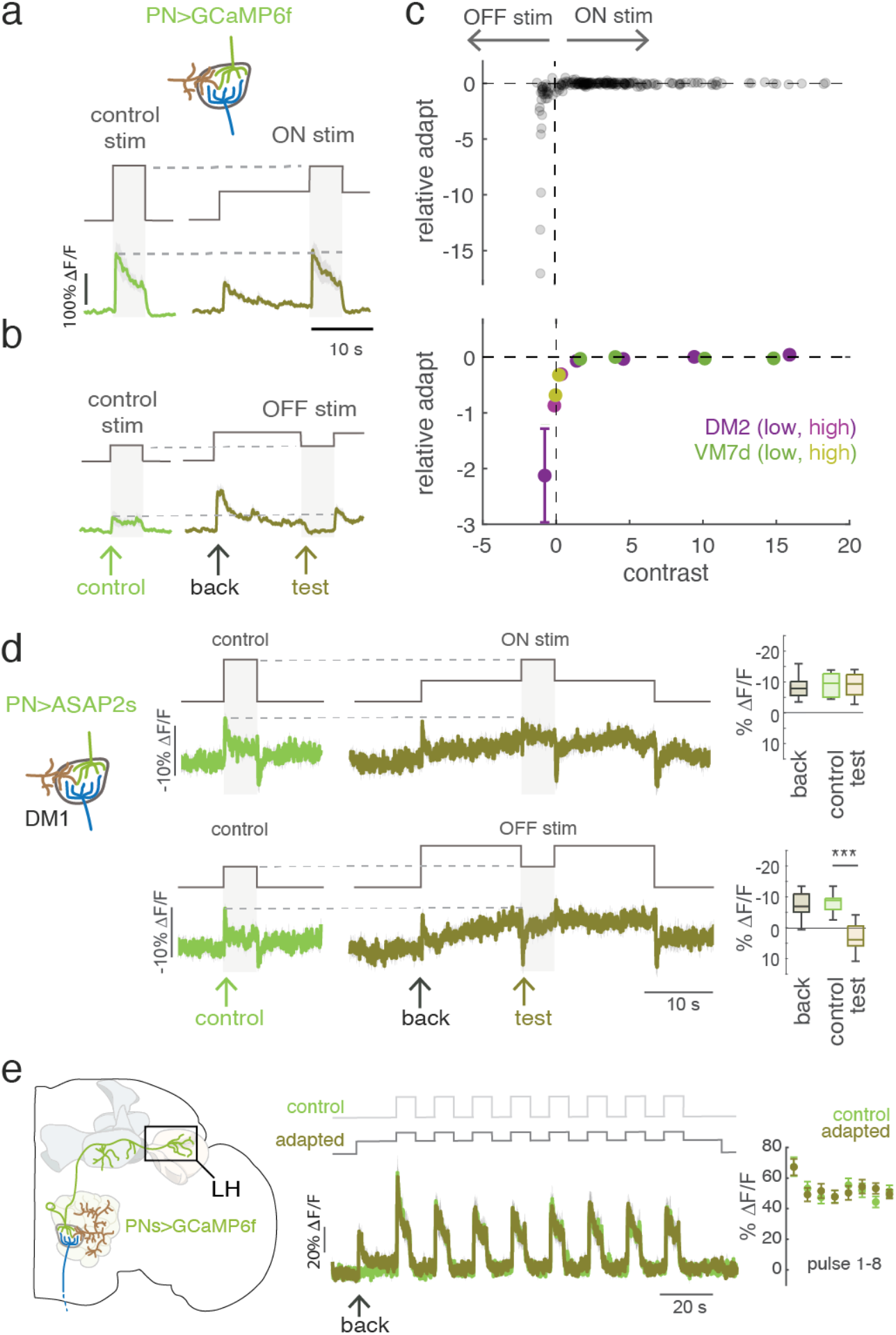
Asymmetric background-invariance of PN responses. **a, b)** Schematics of the stimulus protocol and calcium responses of PNs withing glomerulus DM2 for two examples of ON and OFF stimuli. Colored lines indicate the mean ΔF/F, shaded area SE, N=9-11. The arrows indicate the control (F_control_, green), background (F_back_, black) and test responses (F_test_, olive) used to define the contrast stimulus and relative adaptation. **c)** *Top:* Relative adaptation as a function of contrast for each single trial, including two independent datasets recorded from two different glomeruli. Relative adaptation is defined as (F_test_-F_control_)/F_control_. Since stimulus concentrations are too low to be measured, we use response contrast as a proxy for stimulus contrast (F_test_-F_back_)/F_back_ under the assumption that the response is monotonic. *Bottom*: relative adaptation averaged in bins of contrast; color coded by glomerulus type. *Low* and *high* indicate two different datasets where different ranges of background concentration where used (N=9-11, see also **Fig. S4**). **d)** Control and background adapted responses measure in DM1 using the voltage sensor ASAP2s. Thick lines indicate the mean ΔF/F and the shaded areas SE. Liquid dilution of methyl acetate is 10^−2^, flow dilutions are B6C9 for ON stimuli and B9C6 for OFF stimuli, N=10. Boxplots represent median, quartiles, min/max values and outliers **e)** Calcium responses measured from the PNs axons in the LH, N=6. Odor stimuli were chosen to excite a single glomerulus (DM4, B5C5 with 10^−4^ and 10^−6^ liquid dilutions).

To validate the generality of this observation, we explored a range of ON and OFF stimuli, i.e. odor pulses higher or lower than the adapting background stimulus (**Fig. 6a, b**). PN responses to ON stimuli had similar amplitudes independently of background adaptation (**Fig. 6a, Fig. S4a**), also for high background concentrations (**Fig. S4b)**. In contrast, OFF stimuli, even of small amplitude, always produced large decreases in calcium (**Fig. 6b, Fig. S4b, c)**. To capture the transition between ON and OFF responses irrespective of experimental noise in the generation of the stimuli, we quantified relative adaptation as a function of contrast. We use response *contrast* (defined as the relative difference between the response to the control pulse and the response to the background) as a proxy for stimulus contrast and define *relative adaptation* as the relative difference in the response to the pulse between control and adapted conditions (**Fig. S4c)**. Zero contrast means that the tested stimulus equals the background, and zero adaptation means that PNs respond in a background-invariant manner. By binning the single trials based on contrast value, one can capture the sharp transition between ON and OFF stimuli (**Fig. 6c**), which demonstrates background-invariance only for ON stimuli in the PN dendrites. Responses to OFF stimuli instead fall much below the non-adapted response also for very small contrast values. Therefore the asymmetric background invariance of ORN axonal calcium is preserved in PNs. These observations contrast with a previous study that described background adaptation in PNs firing rates^38^. This could be attributed to a more precise control of test and background stimuli in our experimental setup (**Fig. S4f, g**). However, it could be also due to the different measures of neural activity: calcium versus voltage. PNs’ subthreshold potentials and firing rates follow a linear relationship^38^, therefore, to support our observations, we performed optical voltage recordings from their dendrites. Using the voltage sensor ASAP2s^39^, we show that PN dendrites depolarize similarly (**Fig. 6d**, DM1) or even slightly more (rather than less, **Fig. S4d, e**, DM2 and VM7d) when odor stimuli are presented on a background (ON stimuli). Consistently with calcium measurements, OFF responses are not background invariant (**Fig. 6d**). Finally, we asked whether these properties are preserved in the PNs output. Calcium imaging from the axonal innervations of PNs in the LH shows phasic and background invariant odor responses (**Fig. 6e**) further supporting the relevance of these dynamics for information transmission.

We conclude that both PNs’ dendritic and axonal activities encode a background-invariant representation of ON odor stimuli overcoming the background adaptation of ORN firing rates^13,14^ and the depression of ORN-PN synapses^31,36^.

### PN’s background invariance requires modulation of ORN synaptic release

Next, we asked how the depressing ORN-PN synapses could preserve background invariance. One key difference between presynaptic ORN and postsynaptic PN calcium is their dynamics: tonic in ORNs and phasic in PNs. We reasoned that tonic levels of presynaptic calcium – induced by the sustained background - might be associated with enhanced engagement of the neurotransmitter release machinery (**Fig. 7a**). Conserved Unc13 proteins mediate several types of presynaptic plasticity (e.g. short-term facilitation and fast and long-term potentiation)^40–47^. The two Unc13 splice variants, Unc13A and Unc13B, are differentially organized at the ORN-PN synapses and mediate different postsynaptic currents ^48^. Unc13-dependent plasticity has been linked to three domains that can be modulated directly or indirectly by calcium: the Ca^2+^/calmodulin binding CaM domain (only present in Unc13A), the C1 domain binding diacylglycerol (whose metabolism is Ca^2+^-dependent), and the C2B domain, that binds phosphoinositides in a Ca^2+^-dependent manner^40,49^. The C1 and C2B domain were shown to exert autoinhibition on the vesicle priming reaction, which makes vesicles responsive to action potentials^50^. Activation of the Unc13-C1 domain enhances action-potential induced transmitter release and diminishes multiple forms of presynaptic plasticity (i.e. short-term facilitation, post-tetanic potentiation, and homeostatic potentiation)^47,51–53^. A single amino acid exchange (H→K) in the C1 domain reconstitutes this enhanced potentiation and blocks synaptic plasticity (short term facilitation and potentiation) ^47,52,54^. We used the Unc13-C1 ^HK^ mutation to test if ORN presynaptic plasticity is required to achieve background invariance. We used three stimulus conditions combining two concentrations of the background and test stimulus which, in control flies, led to background-invariant responses in both DL5 ORNs and PNs (**Fig. 7b, c**). It should be noted that in this glomerulus the background compensation is initially overestimated, and background invariance is achieved only at the second odor pulse. Such invariance was lost in Unc13-C1^HK^ mutants for two of the three stimulus conditions tested (**Fig. 7d**). We did not see major effects of the C1 domain mutation for the high contrast condition, which indicates that the calcium modulation probably involves more than one mechanism. Comparing control and mutant flies across three different glomeruli (including DL1 and DM4), we observed significant effects of the mutation on relative adaptation for most stimulus conditions tested (**Fig. 7e**). Unlike expected from a loss of function of Unc13 in ORNs^48^, the C1 gain-of-function mutation had no significant effects on the controls and specifically affected only the response to background adapted stimuli (**Fig. 7f**). This indicates that only activity-dependent modulation of the domain is disrupted, while synaptic transmission itself is not impaired or attenuated in this context, consisted with enhanced, but non-plastic synaptic transmission at the larval NMJ^47^.

**Figure 7.**
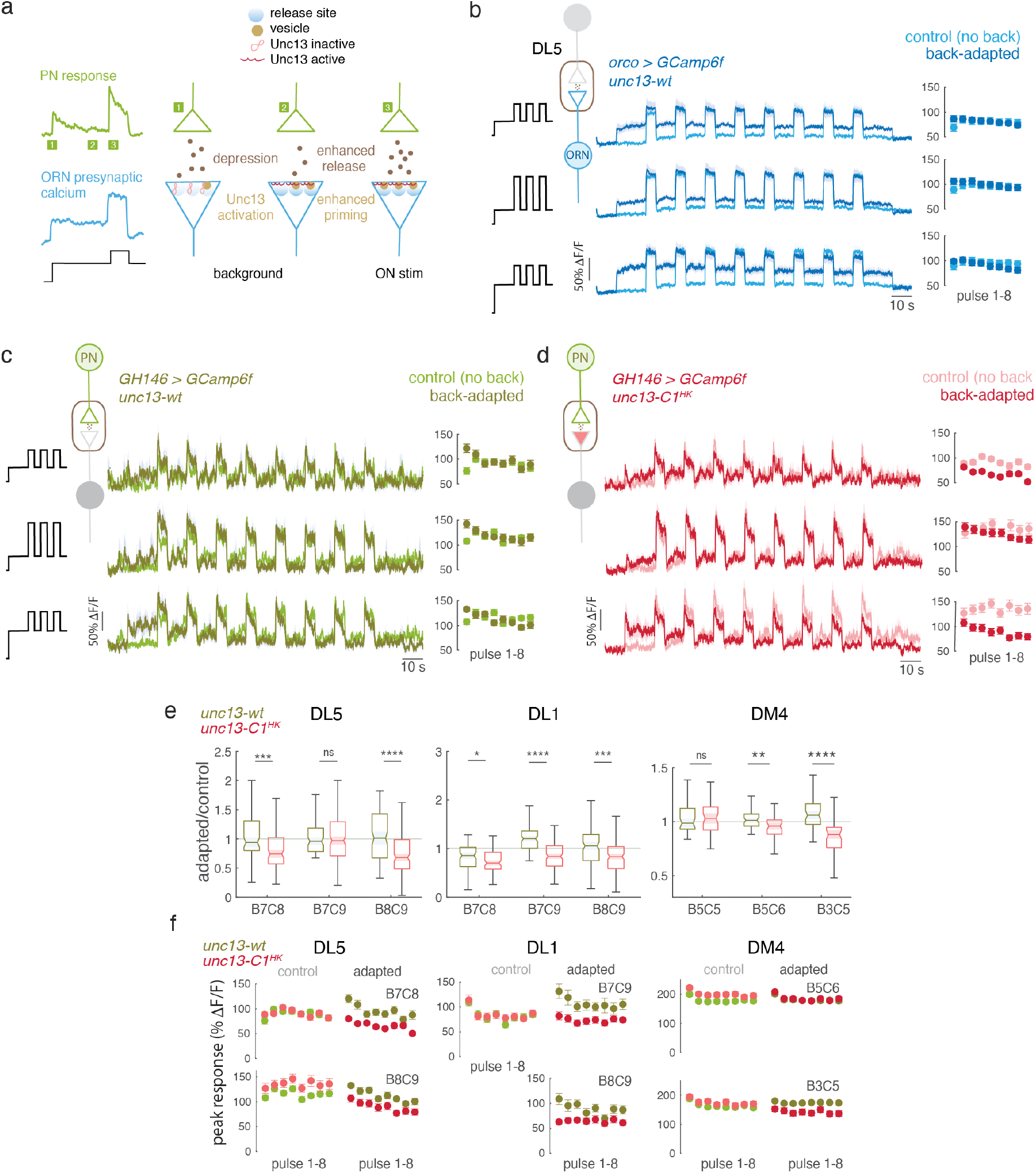
Unc13-C1 domain is required for background-invariant PN response. **a)** Schematics of hypothesis. **b)** Mean calcium response from ORN axon terminal to 8 odor pulses in control and adapted conditions for three combinations of background and test stimuli. Shaded areas indicate SE. Right, quantification of the mean response to each odor pulse. **c**,**d)** same as **b)** for PNs in control and Unc13-C1^HK^ mutant flies (WT N=6-7, mutant N=7-11). **e)** Background adapted response divided by the control for each stimulus condition tested in three different glomeruli. For DL5 and DL1, pure 2-butanone was used for background and pulse. For DM4, we used methyl acetate with liquid dilution of 10^−6^,10^−4^ (B5C5 and B5C6) and 10^−4^,10^−4^ (B3C5). Bar plots are generated from all flies measured and all 8 puffs in each trial (DL1: control N= 6-7, mutant N=7-11, DM4: control N= 6-8, mutant N=6-7). Bar plots indicate median, notch, quartiles, max and min. BXCX indicates the flow rate for background (B) and test (C) stimuli, see Table S3. **f)** Mean peak response for control and Unc13-C1^HK^ mutant flies for 8 consecutives odor puffs in control adapted conditions. Error bars indicate SE.

Overall, we conclude that modulation of Unc13 proteins is necessary to keep PN postsynaptic response background invariant and stable over time. This pathway might act redundantly or synergistically with other calcium-dependent modulations of presynaptic function.

### A population code with asymmetric background invariance

Having identified the mechanisms that ensure background invariance in PNs, we ask how odor information is represented in the background-invariant population responses of PNs compared to the background-dependent population rates of ORNs. We model ORNs based on their peripheral firing dynamics using the amplitude-shift model of adaptation (as in **Fig. 1i)** with the update rule that ensures background invariance (**Fig. 5d** and **Eq. 8** in Methods). We model individual PNs as background invariant and using gaussian noise to recapitulate their sharp response transition between ON and OFF stimuli (**Fig. 6c**) which we attribute to low signal-to-noise in their input (**Fig. S5** and Methods). As shown in **Fig. 1** for ON stimuli, a population of ORNs encodes both intensity and contrast information, but the odor representations shrink close to zero contrast leading to a continuous transition to OFF stimuli (**Fig. 8a, b**). On the contrary, the response space of the PN population exhibits a discontinuity that separates ON and OFF stimuli across different backgrounds while preserving a background-invariant representation of ON stimuli (**Fig 8c**).

**Figure 8.**
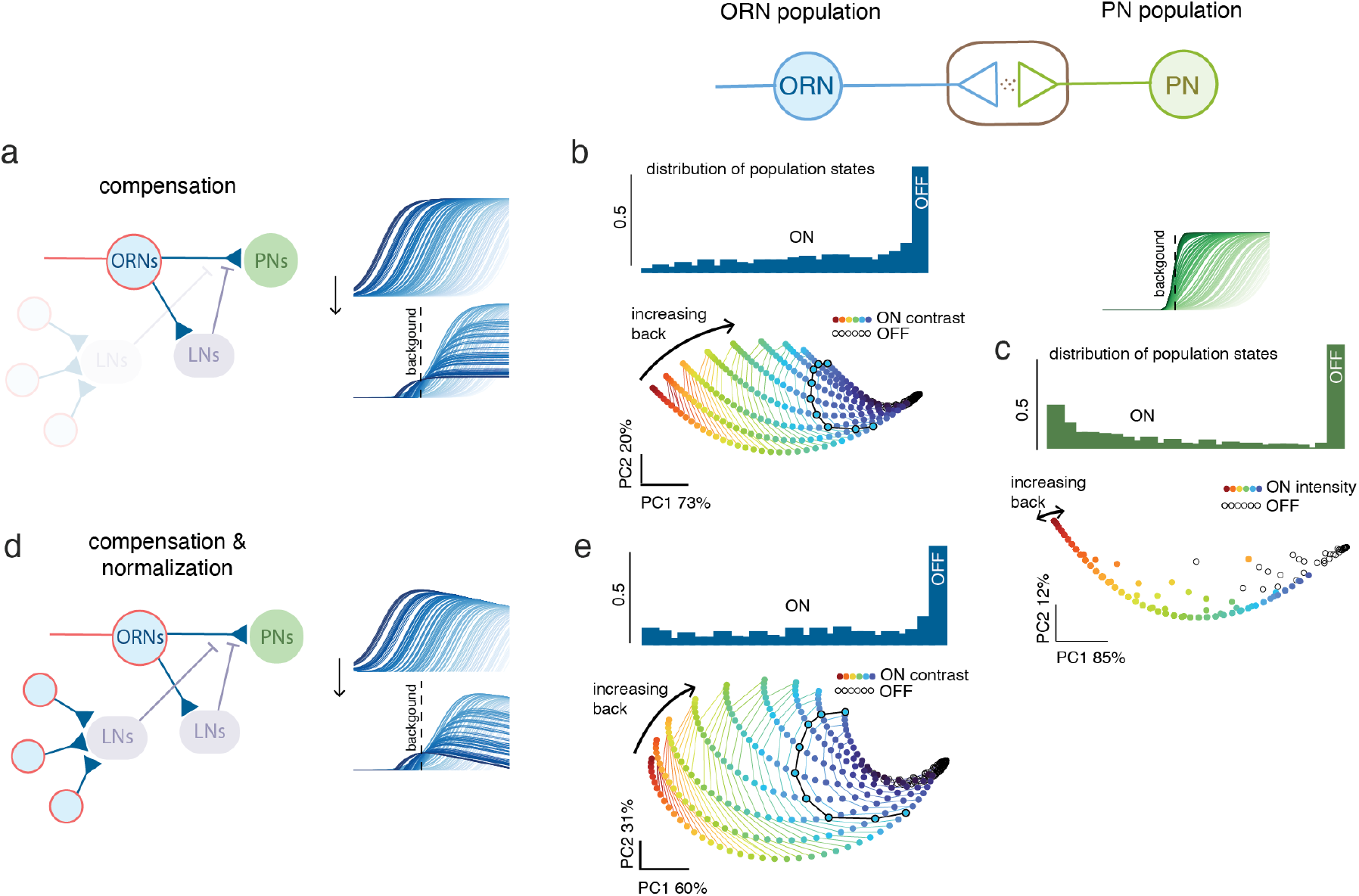
Combinatorial representation of ON and OFF stimuli in ORN and PN populations: **a)** Schematics of the circuit model that implements synaptic compensation of firing rate adaptation. *Right*: firing response curves for a population of ORNs with different stimulus sensitivities in control (*top*) and background adapted conditions (*bottom*). **b)** 2D-manifold of the population response to stimuli of different ON (color coded) and OFF (black) contrast in different background adapted conditions. The histogram represents the distribution of population states calculated on the first principal component. **c)** Same as **b)** for PNs, but responses are color-coded by stimulus intensity. PNs ON background-invariance was modeled as described in **Fig. S4. d)** Schematics of the circuit model that implements both synaptic compensation and interglomerular normalization. *Right*: response curves for a population of ORNs with different stimulus sensitivities in control (top) and background adapted conditions (bottom). **e)** Same as **b)** but responses of single neurons are normalized by a term proportional to stimulus contrast.

Since the background-invariant responses of PNs do not encode contrast explicitly at the population level, there is a remaining question of why ORN populations should encode this feature. We have shown both experimentally (**Fig. 3**) and computationally (**Fig. 5**) that firing rate compensation can be performed in a glomerulus-intrinsic manner. However, when multiple glomeruli are active, background invariance must be achieved in a coordinated way across the AL. We ask whether this task could be achieved by multiglomerular LNs that would be able to read out both intensity and contrast information from ORN activity (**Fig. 1**). LNs mediate a form of normalization that relaxes the non-linearity of the ORN-PN transformation^32,33^. This normalization is mostly presynaptic^33,55^ and relies on a measure of overall activity across receptor types. We studied the effects of such normalization by including an interglomerular LN to the local ORN-PN circuit (**Fig. 8d** and Methods). Adding this interglomerular normalization factor enhances the encoding of low contrast stimuli in ORNs (**Fig. 8e**). We show analytically that with this normative input at the presynapses, ON background invariance of the PNs response can only be achieved by the amplitude-shift mechanism if multiglomerular LNs encode contrast information, in agreement with our hypothesis (Methods). Remarkably, in the absence of background stimuli, the encoding of contrast equals the encoding of intensity consistently with reported normative function of the LNs^33^. Using a divisive rather than a linear model of lateral inhibition leads to similar results (Supplementary Material, **Appendix 1** and **Fig. S3**).

Overall, we propose that the AL implements a circuit computation that extracts contrast information encoded in ORNs to enhance the discriminability between ON and OFF stimuli in postsynaptic PNs while preserving information of odor identity for ON stimuli.

## Discussion

Odors are detected by single sensory neurons that work combinatorially to represent complex natural stimuli at the population level. Despite the key role of adaptation for information coding in both single neurons and neuronal populations, the adaptive properties of the olfactory system downstream of the receptors have been largely overlooked. In this work, we investigated adaptation as a coding strategy of single ORNs that shapes the encoding capabilities of olfactory neural populations. By combining experiments and computational models, we identified a novel property of the olfactory pathway that we defined as asymmetric background invariance. We show that the adapted firing responses of single ORNs are compensated to achieve an asymmetric representation of ON and OFF contrast stimuli, where ON responses are background invariant while OFF responses collapse in the representation space of the downstream populations, the PNs. Mechanistically background invariance requires a feedback compensation induced by ORN synaptic release and involves a GABAergic inhibitory pathway that modulates ORN presynapses. We propose that this feedback compensation facilitates vesicle release through interaction with the Unc13-C1 domain, which stabilizes the background invariance of the PNs. The PN population therefore preserves ON stimulus identity necessary for downstream computations.

### The role of adaptation at the single neuron and population level

Efficient coding principles provide an explanation for how single sensory neurons adapt their response function to changes in stimulus statistics over behaviorally^2,9,10^ or evolutionary^56–58^ relevant time scales. Similar principles have been used to understand how the response function of neurons in a population should adjust to tile the stimulus space in a flexible manner^59^. This is of particular importance to understand coding principles in the olfactory system, which uses a repertoire of receptors to detect and discriminate odors within neuronal populations. Receptor repertoires are tuned, through evolutionary times, to the ecology of the animal^60^, leading to efficient sensory codes in the ecologically relevant stimulus space^61^. A similar adaptive strategy also applies on shorter timescales. For instance, the expression levels of the receptors are plastic in mammals, providing a mechanism to optimize the composition of a receptor repertoire to changes in the environment^62^.

Similar adaptive changes in the coding capability of a receptor repertoire could in principle occur also on shorter time scales, but there is limited evidence of such a strategy in the olfactory system^28^. In this work, we provided evidence that the fly olfactory system does not adapt the odor representation to the background. Rather, it implements mechanisms that keep odor representations robust with respect to changes in the background. While efficient coding theories of single neurons predict the conventional sensitivity-shift strategy widely observed in the visual system, both ORNs and PNs do not follow this principle. ORNs adapt by modifying their response amplitude (and not their sensitivity), which ensures the compensation necessary to achieve ON background invariance in downstream neurons, the PNs. Therefore, even though a persistent stimulus (the background) induces synaptic depression and a corresponding decrease in postsynaptic activity^36^, it also primes the synapses to respond to ON stimuli as strongly as in the absence of background. The mechanisms underlying adaptation involve inhibitory feedback potentially provided by LNs. Both dynamic and stationary models for ORN adaptation through inhibitory feedback confirm the role of an amplitude-based adaptation that can come as a single amplitude shift or as a combination of both amplitude-shift and -compression.

Our findings raise the question of why ORNs would adapt at all if the goal was to encode absolute stimulus intensities. We argue that firing rate adaptation is beneficial to the asymmetric encoding of ON and OFF stimuli, where only ON stimuli preserve stimulus identity, while OFF stimuli collapse into a low-response state. Such asymmetry would not be possible if the ORNs simply encoded stimulus intensity. From a more speculative perspective, another potential benefit of adapting firing rate and then compensating it downstream could be a lower total energetic cost of sending information through the brain with no loss. However, further work is necessary to assess whether the synaptic computations that compensate firing rate adaptation are indeed less energetically costly than implementing a non-adaptive spiking code.

### The role of asymmetric odor encoding downstream of PNs

PNs send odor information to both the LH and MB, contributing to higher order computations. Odor representations in the Kenyon cells (KCs) of the MB are used to form associative memories based on the co-occurrence of an odor and a reinforcement^63^. For memory retrieval, it is critical that the same KCs are activated by the conditioned stimulus in training and test phase and therefore it is critical that the odor is represented independently of possible backgrounds. Altogether, memory retrieval provides a potential function for background-invariant encoding in PNs. Consistently with this hypothesis, a recent study showed that locusts learn in a background-invariant manner^64^, suggesting that the MB has access to a background-invariant representation of odors.

Odor cues are used not only to evaluate, but also to explore the environment. Animals use odor gradients^65,66^, plume statistics^67,68^ and odor motion^69^ to direct their behavior in complex olfactory environments. These computations require the detection of gradients and contrast in an odor-specific manner. It is plausible that such computations do not occur in the AL, where odor representations are formed, but in downstream neurons that can read out a population response from the AL to then compute stimulus-specific information on contrast. The LH provides the cellular complexity^70,71^ to perform odor-specific computations in parallel^72^, including the encoding of contrast^73^. We suggest that the encoding of background-invariant ON responses through the compensation of ORNs adaptation is essential for downstream stimulus-specific computations that drive key behaviors in flies. These computational requirements do not generalize to OFF stimuli, which have a very different relevance for the fly behavior compared to ON stimuli, leading to an explorative search rather than an oriented behavior ^65,74^.

### Cell intrinsic and circuit mechanisms that maintain ON stimulus invariance

Asymmetric background invariance is the most robust observation we made across all our measurements in both ORNs axons and PNs dendrites. However, we also encountered differences across glomeruli. Firstly, ORN calcium dynamics range from being slightly phasic (DM1), completely flat (DL1), to slightly ramping up (DL5). This seems to correlate with background invariance to be achieved already on the first (DM1), or only on the second or third (DL1, DL5) odor pulse. Moreover, we showed that background invariance of DM1 and DL1 ORNs requires synaptic transmission, whereas DL5 background invariance does not. Moreover, blocking GABA receptors has differential effects across glomeruli, which unlikely can be attributed solely to differences in drug penetration. Glomeruli such as DL5, which do not receive substantial GABAergic inputs under the condition tested, may instead rely on cell-intrinsic mechanisms - for example the regulation of internal calcium storages^75,76^. In other glomeruli, these intrinsic mechanisms may act in combination with feedback inhibition, which is recruited in a stimulus-dependent manner.

In all tested glomeruli, we find that a gain-of-function mutation of the Unc13-C1 domain, Unc13-C1^HK^, occludes PNs background invariance. So far, we do not have direct evidence that plasticity involving Unc13 is mediated by the tonic and background-invariant presynaptic calcium nor that it is downstream of lateral inhibition. However, our current working model postulates a link between presynaptic calcium regulation and enhanced vesicle priming. As the Unc13-C1^HK^ mutant does not abolish background invariance for all stimulus conditions, other calcium-dependent mechanisms are likely involved in this plasticity, perhaps through modulation of other synaptic proteins (e.g. synaptotagmin-3 or -7^77^).

Overall, our data suggest a key role for presynaptic plasticity in PN background invariance, linking the same molecular pathways usually studied under artificially induced homeostasis^40,47,78^ to sensory computations in natural conditions. At ORN-PN synapses, the homeostatic set point is the non-adapted response. Differently from other models of synaptic homeostasis, the feedback compensation is not calculated based on synaptic output (PN response) but based on the input drive to synaptic release (ORN firing rate). This synaptic strategy is, to our knowledge, unique in its design and perhaps specific to the computational demands of the olfactory system.

### The undoing of a neural computation: evolutionary considerations

Undoing an upstream computation is rather an uncommon observation that becomes counterintuitive from an efficient coding perspective, where computations are expected to optimize information processing across neuronal layers. However, if we look at neural circuits as ongoing projects of evolution that continue to adapt to optimize behaviorally relevant trade-offs^79,80^, more examples of undoing can be found. For example, some amacrine cells in the visual system undo the color opponency calculated in upstream neurons to preserve the balance between opponent and non-opponent units in the retina^81^ such that both types of information are transferred in downstream neurons. The logic of this computation could result from the order in which these mechanisms evolved (discussed in^81^).

From an evolutionary perspective, the olfactory system is designed to ensure adaptability, which is implemented primarily by the evolution of chemosensory genes. Our considerations focused on the combinatorial aspect of odor coding, and therefore the fact that stimuli are encoded in populations of sensory neurons, with identity defined by the OR they express. However, this system evolved from an ancestor that expressed only one or few ORs^82^. In such a low dimensional system, firing rate adaptation might not only have been parsimonious, but also an efficient strategy for odor encoding, as it is the case for visual neurons. The olfactory system might have selected for mechanisms that support background invariance only after the emergence of a combinatorial coding strategy with multiple ORs.

Olfactory systems also evolved specialized channels that encode unique chemicals, such as pheromones^83^ or harmful substrates^84^, crucial in certain ecological contexts. These stimuli therefore do not rely on a population code and might implement different strategies for adaptation. A combination of mechanisms can be therefore expected within the same sensory system depending on the function of specific subcircuits. Further insight into these questions might benefit from the study of downstream areas like the LH that read out combinatorial or specialized representations for behavioral-relevant computations.

## Material and methods

### Experimental model / fly husbandry

All flies were kept on a standard molasses-based food at 25°C and on a 12h:12h light-dark cycle or at room temperature in the laboratory. For optogenetic experiments, flies were kept in the dark and on food that was supplement with 1mM all-trans retinal (Sigma-Aldrich, Table S1) for at least 4 days before experiments. For shibire^ts^ experiments, flies were incubated for at least 1h at 36-37°C before the start of recordings, which lasted a maximum of 45min. Female flies were used for all experiments. The complete genotypes are given in Table S2.

### Fly mutagenesis Unc13-C1^HK^

The mutation of the Unc13-C1 domain was based on the corresponding mutation of the mouse homologues Munc-13-1 where histidine 567 was changed to lysine^52^. Sequence alignment with *Drosophila* Unc13A identified histidine 1723 as the relevant target. Note that the two *Drosophila* splice isoforms Unc13A and Unc13B harbor a C1 domain^46^ and that this modification therefore affects both. CRISPR-mediated mutagenesis was performed by WellGenetics Inc. using modified methods of Kondo and Ueda^85^ to generate a break in unc-13/CG2999. A PBacDsRed cassette containing two PBac terminals and 3xP3-DsRed and two homology arms with point mutations (one to induce the amino acid change and one to introduce a TTAA sequence to recognize proper integration on the other homology arm) were used for repair. The DsRed marker was intermittently used to identify flies with successful integration events in the F1 generation. The marker was later excised and the presence of the point mutation and successful removal of the cassette in the final fly line validated by PCR and sequencing from genetic DNA.

### *In vivo* calcium imaging

Flies 6-10 days post-eclosion were anesthetized on ice and mounted on a custom holder using the Bondic repair system (Bondic cartridge with Bondic liquid plastic and Bondic UV-LED). The proboscis was secured, and the maxillary palps were covered using low melting temperature wax. Saline solution (5mM Hepes, 130 mM NaCl, 5mM KCl, 2 mM MgCl2, 2mM CaCl2, 36 mM Saccharose – pH 7.3) was added and the cuticle covering the fly’s brain, as well as obstructing trachea and fat tissue, were removed.

Functional imaging was performed on an Investigator two-photon microscope (Bruker) coupled to a tunable laser (Spectraphysics Insight DS+) with a 25×/1.1 water-immersion objective (Nikon). Laser excitation was tuned to 920 nm, and less than 20 mW of excitation was delivered to the specimen. Emitted light passed through a SP680 short-pass filter, a 560 lpxr dichroic filter and a 525/70 filter. PMT gain was set to 850 V. The microscope was controlled with the PrairieView (5.4) software.

### Stimulation

Test and control stimuli were 5 sec long, background stimuli were 15 sec. In Fig. 4,5,6, test and control stimuli were repeated 8 times with 10 sec between repetitions.

#### Optogenetics

light from a 625 nm LED was directed to the fly’s antenna using an optic fiber. The LED was controlled by the imaging software and activated during the laser flight-back, allowing simultaneous acquisition and excitation. In electrophysiological experiments, the LED was controlled by an Arduino board.

#### Odor delivery

a flow of clean air was presented to the fly continuously (1L/min). To this main airflow, either an odor (odorant airflow; 100mL/min) or more clean air (balancer airflow; 100 mL/min) was added through a solenoid valve (LEE), so that the final airflow reaching the fly was always around 1.1L/min. To create the gas dilutions for the odorants, four mass flow controllers (MFCs) were used (Analyt-MTC) and controlled using a custom MATLAB (MathWorks) script and an Arduino board. Another 3 MFCs were required to keep the constant airflows for the main airflow and the balancer airflows. Odors were prepared in 20ml glass vials as a liquid volumetric dilution with a total volume of 5 mL in mineral oil. Airflow dilutions used are described in Table S3. Liquid dilutions were prepared in 20 ml glass vials always to a total volume of 5 mL in mineral oil.

### Electrophysiology

Single-sensillum recordings were performed as previously described^14^ using a silver-chloride electrode and glass pipettes filled with sensillum lymph ringer. Electrical signals were amplified using an extracellular amplifier (EXT-02F-1, npi) with head stage (EXTEH), bandpass filtered (300–5000 Hz), digitized at 20KHz using a NI board (NI-6212). Data were acquired with the MATLAB toolbox kontroller^27^ (https://github.com/emonetlab/kontroller). Spikes were sorted using a custom MATLAB script. Odor delivery: flies were exposed to a constant airflow (1L/min) and an odor stimulus was delivered by switching a 3-way solenoid valve that directed a secondary airflow (100 mL/min) through a Pasteur pipette as in^14,36^. The pipette contained a filter paper with 50ml of the odor dilution. Volumetric odor dilutions were prepared in either mineral oil or MiliQ Water. Stimuli were controlled by custom-made software in MATLAB and Arduino.

### Pharmacology

Acetylcholine receptor blockage was performed using 100 μM atropine + 20 μM tubocurarine + 200 nM imidacloprid (IMI). Stock solutions of 2 mM atropine and 20 mM tubocurarine were prepared in water and aliquoted and frozen at −20°C. Stock solution of 1 mM IMI was prepared in DMSO and kept at room temperature. Final solution was dissolved in saline solution used for *in vivo* calcium imaging and kept at 4 °C. For controls, instead of the drug solution, vehicle solution was applied, consisting of the saline solution previously mentioned with 5.1% water + 0.02% DMSO. It was also kept at 4 °C. GABA receptor blockage was done using 25 μM CGP54626 + 5 μM picrotoxin (PTX). Stock solutions of 25 mM CGP54626 and 100 mM PTX were prepared in DMSO, aliquoted and kept at −20°C. Final solution was dissolved in saline solution used for *in vivo* calcium imaging and kept at 4 °C.

For all pharmacology experiments, flies were prepared for *in vivo* calcium imaging as explained before, with the addition of a final step during dissection where the brain was de-sheathed. All flies were imaged before application of drug solutions with the same stimulation protocol used after drug was added in bath application.

For acetylcholine receptor blockage experiments, normal saline solution was replaced by 100 μM atropine + 20 μM tubocurarine + 200 nM IMI solution 2 consecutive times and then again 10 min after.

Recordings were started 20 min after the drug solution was initially applied. Vehicle solution was applied in the same manner to control flies.

For GABA receptor blockage, normal saline solution was replaced by 25 μM CGP54626 + 5 μM PTX. This was repeated 1 min, 2 min and 10 min after this initial drug application. Recordings were started 40 min after the drug solution was initially applied.

### Adaptation model: population response

For a population of ORNs, we model the peak firing rate of each individual neuron *i, F*_i_(*s*), as a function of the odor stimulus, *s*, as

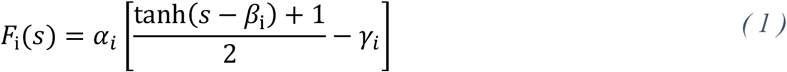

where we have included an explicit dependence on the parameters *α*_*i*_, *β*_i_ and *γ*_*i*_ that control the amplitude-compression, sensitivity-shift, and amplitude-shift adaptation respectively. Neurons within the population are set with an initial random stimulus tuning *β*_i_ and equal parameters α_*i*_ = α and *γ*_*i*_ = *γ*. The adapted firing rate, *F*′_i_(*s*), is described by the same equation but different parameters, *β*′_i_, *α*′_*i*_ and *γ*′_*i*_, which are updated in a background response-dependent way. For the simulations in Fig. 1, we used the update rules,

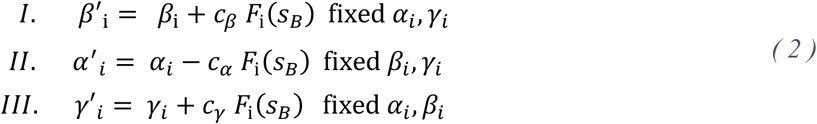

with *c*_*α*_ = *c*_*γ*_ =0.8 and *c*_*β*_ = 1. Our results in Fig 1 show that the adaptation mechanisms II and III (response domain) lead to a population code whose variance is mostly explained by two main principal components. Yet, for the mechanism I (sensitivity domain), only one component is needed. We tested the generality of these results for a larger set of weight parameters *c*_*β*_, *c*_*α*_, *c*_*γ*_ ∈ {0.5,0.6, …, 1.5} for which we quantified the ratio of explained variance between the two first principal components. For each adaptation mechanism we got the ratios *r* _*β*_ = 15.46 ± 0.76, *r*_*α*_ = 1.62 ±0.53 and *r*_*γ*_ = 1.63 ± 0.51, where the error corresponds to the std across all the different weight parameters. Ratios near to unity indicate that both principal components contribute significantly to the population response, while large ratio values indicate a predominant representation of the first principal component. Altogether, we show that the results shown in Fig. 1 hold for different weights in the update rules of Eq. (1) and Eq. (2).

### Adaptation model: firing rate and calcium response

We model the ORN calcium response *R* as a linear function of the difference between the ORN firing rate, *F*_*E*_(*s,t*),and an inhibitory rate, *F*_*I*_(*s,t*),coming from LNs:

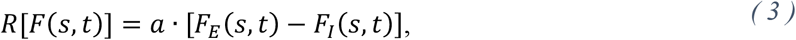

where *a* is the is the proportionality factor between calcium and firing rate in absence of background as shown in **Fig. 2e**. We model the dynamics of each ORN firing rate as the exponential process:

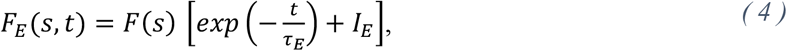

where τ_*E*_ is the excitatory time constant, *F*(*s*) is the firing rate described by Eq. (1) and *I*_*E*_ is the excitatory baseline. We consider the inhibitory drive to have similar dynamics, inherited from the ORN responses, but a different baseline, that is τ_*I*_ = τ_*E*_ and *I*_*E*_ – *I*_*I*_ = Δ_*I*_ = 0.05 respectively. Note that here we do not model the LN firing dynamics, but a general inhibitory input to the pre-synapse. While some LNs types have been shown to have more transient activity^86^, this inhibitory drive could result from specific subtypes or from the signaling cascade downstream of the receptors^87^.

Using Eq. (4) under these excitatory and inhibitory response conditions, the calcium response of Eq. (3) becomes proportional to the difference between the excitatory and inhibitory rates, that is,

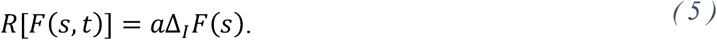

The calcium sustained response to a given stimulus *s* in the presence of a background, *s*_*B*_, is proportional to the sum of the sustained calcium response to the background and the stimulus calcium response after the firing rate is adapted, that is,

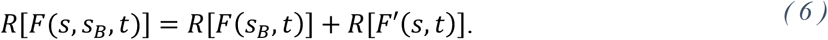

The background invariance of ORN calcium responses implies that *R*[*F*(*s, s*_*B*_,*t*)] = *R*[*F*(*s,t*)] for any background. By using Eq. (5) in Eq. (6) we obtain that the adapted firing rate response *F*′(*s*) has to satisfy:

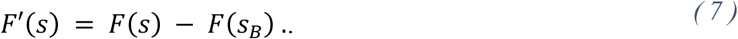

This implies that firing rate adaptation has to occur as a change in the response domain rather than in the stimulus domain. Using the firing rate model defined in Eq. (1), we calculate the adaptation rules that satisfy this calcium invariance for each type of adaptation strategy. Specifically, we calculate the adapted value of the parameters that control each of the three adaptation mechanisms considered:

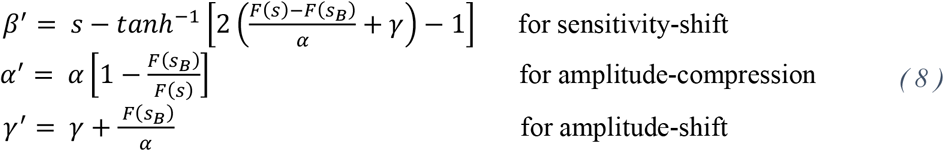

This demonstrate that the only adaptation mechanism that depends uniquely on the background adaptation response and satisfies the calcium invariant condition is the amplitude-shift adaptation. The other two mechanisms require information of both the adapted and non-adapted responses, which are not accessible simultaneously to the circuit.

### Adaptation model: effects of inter-glomerular activity

To understand how inter-glomerular activity further shapes the adapted responses of ORNs, we modeled a second inhibitory mechanism that reads out odor information from the population of ORNs and acts at the single ORN level (see Fig. 8). Specifically, we introduce in our model an LN neuron that connects to several glomeruli^32,33^ and provides an inhibitory input to the presynaptic terminals. This can be formalized by modifying Eq. (3) as,

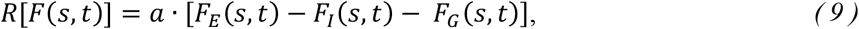

where *F*_*G*_(*s,t*) = ∑_*i*_ *ω*_*i*_ *F*_*i*_(*s,t*) corresponds to the LN term that integrates the ORN population activity. Assuming that the dynamics of the global feedback is slower than the dynamics of the local feedback, our previous arguments hold up to Eq (6). Similarly, as we did before, we use Eq. (9) to write a new relation between the calcium responses of non-adapted and adapted ORNs:

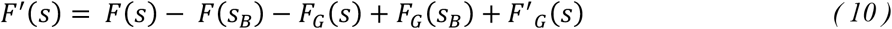

We showed in **Fig. 1** that a population of ORNs encodes both stimulus intensity and contrast in a two-dimensional manifold, making it possible for two downstream neurons to extract such an information as a linear combination of single ORNs. Consequently, if we assume that the LN neuron that connects to multiple glomeruli extracts contrast information from the population, we can write *F*_*G*_(*s*) = *s*−*s*_*B*_, which at zero background equals the stimulus intensity. We can then write Eq. (10) as,

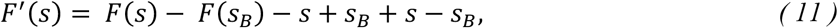

showing that the amplitude-shift mechanism continues to support ON-background invariance if contrast information is extracted from ORN populations by inter-glomerular LNs. In the absence of an odor background, the encoding of contrast information equals the encoding of the intensity value, which is in agreement with the literature on the role of inhibitory LNs^32,33,55^.

### PN model of asymmetric background invariance

We show that PNs inherit the ON background-invariant responses from ORN calcium responses. Consequently, we model the adapted responses of PNs in the same way as for the calcium responses of ORNs (Fig. 5). To account for the sharp but continuous transition from OFF to ON responses near small contrast stimuli (Fig 6c), we implement a noise model for the relative adapted responses of PN neurons with a probability N(*s*−*s*_*B*_, *σ*), where *s*_*B*_ is the background stimulus and *σ* is the noise standard deviation. We show that this noise model reproduces the transition from OFF to ON stimuli observed in the experiments (See S4). We then use this model to simulate the adapted responses of PNs (Fig. 8c).

## Data Availability

All data are available in the main text or the supplementary materials. Raw data, scripts and software for data curation and analysis have been deposited in Zenodo and are accessible at 10.5281/zenodo.15937307.

## Code Availability

Additional resources are available on GitHub https://gitlab.rlp.net/mrtlllab/brandao_ramirez_adaptation2024

## Acknowledgment

We thank Filippo Calzolari for reading the first version of the manuscript, Giovanni D’Uva for developing methodology, Christopher Schnaitmann for help with *in vivo* imaging and comments on the manuscript, Christian Daniel for help with graphics, members of the Martelli and Silies lab for discussions, Sabine Schmitt, Simone Renner and Jonas Chojetzki for technical and administrative support. We further thank Douglas Storace for initial discussions on this project, Henrike Scholz for sharing fly lines and Mathias Böhme for help in the planning of the mutagenesis of the Unc13-C1 domain.

## Author contribution

Conceptualization: SCB, CM

Methodology: SCB, LR, PZ, AMW, MS, CM

Investigation: SCB, LR, PZ, CM

Formal analysis: LR, CM

Visualization: SCB, LR, PZ, CM

Supervision: CM

Writing—original draft: LR, CM

Writing—review & editing: SCB, LR, AMW, MS, CM

## References

1. Wark, B., Lundstrom, B. & Fairhall, A. Sensory adaptation. Curr Opin Neurobiol 17, 423–429 (2007).

2. Laughlin, S. B. The role of sensory adaptation in the retina. Journal of Experimental Biology 146, 39–62 (1989).

3. Simoncelli, E. P. Vision and the statistics of the visual environment. Curr Opin Neurobiol 13, 144–149 (2003).

4. Brenner, N., Bialek, W. & De Ruyter Van Steveninck, R. Adaptive Rescaling Maximizes Information Transmission. Neuron 26, 695–702 (2000).

5. Laughlin, S. B. & Hardie, R. C. Common strategies for light adaptation in the peripheral visual systems of fly and dragonfly. Journal of Comparative Physiology □ A 128, 319– 340 (1978).

6. Gür, B. et al. Neural pathways and computations that achieve stable contrast processing tuned to natural scenes. Nature Communications 2024 15:1 15, 1–18 (2024).

7. Malnic, B., Hirono, J., Sato, T. & Buck, L. B. Combinatorial receptor codes for odors. Cell (1999) doi:10.1016/S0092-8674(00)80581-4.

8. Hallem, E. a & Carlson, J. R. Coding of odors by a receptor repertoire. Cell 125, 143– 60 (2006).

9. Benda, J. Neural adaptation. Current Biology 31, R110–R116 (2021).

10. Weber, A. I., Krishnamurthy, K. & Fairhall, A. L. Coding Principles in Adaptation. Annu Rev Vis Sci 5, 427–449 (2019).

11. Gutnisky, D. A. & Dragoi, V. Adaptive coding of visual information in neural populations. Nature 2008 452:7184 452, 220–224 (2008).

12. Reisert, J. & Matthews, H. R. Adaptation of the odour-induced response in frog olfactory receptor cells. Journal of Physiology (1999) doi:10.1111/j.1469-7793.1999.0801n.x.

13. Nagel, K. I. & Wilson, R. I. Biophysical mechanisms underlying olfactory receptor neuron dynamics. Nat Neurosci 14, 208–216 (2011).

14. Martelli, C., Carlson, J. R. & Emonet, T. Intensity invariant dynamics and odor-specific latencies in olfactory receptor neuron response. Journal of Neuroscience 33, (2013).

15. Strausfeld, C. Z. & Kaissling, K.-E. Localized adaptation processes in olfactory sensilla of Saturniid moths. Chem Senses 11, 499–512 (1986).

16. Lemon, W. & Getz, W. Temporal resolution of general odor pulses by olfactory sensory neurons in American cockroaches. Journal of Experimental Biology 200, 1809–1819 (1997).

17. Raman, B., Joseph, J., Tang, J. & Stopfer, M. Temporally diverse firing patterns in olfactory receptor neurons underlie spatiotemporal neural codes for odors. Journal of Neuroscience 30, 1994–2006 (2010).

18. Kurahashi, T. & Menini, A. Mechanism of odorant adaptation in the olfactory receptor cell. Nature 385, 725–729 (1997).

19. Dalton, P. Psychophysical and behavioral characteristics of olfactory adaptation. Chem Senses 25, 487–492 (2000).

20. Ramaswami, M. Network plasticity in adaptive filtering and behavioral habituation. Neuron 82, 1216–1229 (2014).

21. Sinding, C. et al. New determinants of olfactory habituation. Sci Rep 7, 41047 (2017).

22. Wilson, D. A. Habituation of odor responses in the rat anterior piriform cortex. J Neurophysiol 79, 1425–1440 (1998).

23. Fishilevich, E. et al. Chemotaxis behavior mediated by single larval olfactory neurons in Drosophila. Current Biology (2005) doi:10.1016/j.cub.2005.11.016.

24. Baker, K. L. et al. Algorithms for Olfactory Search across Species. J. Neurosci. 38, 9383–9389 (2018).

25. Sourjik, V. & Wingreen, N. S. Responding to chemical gradients: bacterial chemotaxis. Curr Opin Cell Biol 24, 262–268 (2012).

26. Louis, M. Mini-brain computations converting dynamic olfactory inputs into orientation behavior. Curr Opin Neurobiol 64, 1–9 (2020).

27. Gorur-Shandilya, S., Demir, M., Long, J., Clark, D. A. & Emonet, T. Olfactory receptor neurons use gain control and complementary kinetics to encode intermittent odorant stimuli. Elife 6, e27670 (2017).

28. Martelli, C. & Storace, D. A. Stimulus Driven Functional Transformations in the Early Olfactory System. Front Cell Neurosci 15, (2021).

29. Brandão, S. C., Silies, M. & Martelli, C. Adaptive temporal processing of odor stimuli. Cell Tissue Res 383, 125–141 (2021).

30. Su, C.-Y., Menuz, K. & Carlson, J. R. Olfactory Perception: Receptors, Cells, and Circuits. Cell 139, 45–59 (2009).

31. Kazama, H. & Wilson, R. I. Homeostatic matching and nonlinear amplification at identified central synapses. Neuron 58, 401–413 (2008).

32. Olsen, S. R. & Wilson, R. I. Lateral presynaptic inhibition mediates gain control in an olfactory circuit. Nature 452, 956–960 (2008).

33. Olsen, S. R., Bhandawat, V. & Wilson, R. I. Divisive normalization in olfactory population codes. Neuron 66, 287–299 (2010).

34. Banerjee, A. et al. An Interglomerular Circuit Gates Glomerular Output and Implements Gain Control in the Mouse Olfactory Bulb. Neuron 87, 193–207 (2015).

35. Laughlin, S. B. & Hardie, R. C. Common strategies for light adaptation in the peripheral visual systems of fly and dragonfly. Journal of Comparative Physiology □ A (1978) doi:10.1007/BF00657606.

36. Martelli, C. & Fiala, A. Slow presynaptic mechanisms that mediate adaptation in the olfactory pathway of Drosophila. Elife 8, (2019).

37. Barth-Maron, A., D’Alessandro, I. & Wilson, R. I. Interactions between specialized gain control mechanisms in olfactory processing. Curr Biol 33, 5109–5120.e7 (2023).

38. Cafaro, J. Multiple sites of adaptation lead to contrast encoding in the Drosophilaolfactory system. Physiol Rep 4, e12762 (2016).

39. Chamberland, S. et al. Fast two-photon imaging of subcellular voltage dynamics in neuronal tissue with genetically encoded indicators. Elife 6, (2017).

40. Jusyte, M. et al. Unc13A dynamically stabilizes vesicle priming at synaptic release sites for short-term facilitation and homeostatic potentiation. Cell Rep 42, (2023).

41. Augustin, I., Rosenmund, C., Südhof, T. C. & Brose, N. Munc13-1 is essential for fusion competence of glutamatergic synaptic vesicles. Nature 400, 457–461 (1999).

42. Walter, A. M., Böhme, M. A. & Sigrist, S. J. Vesicle release site organization at synaptic active zones. Neurosci Res 127, 3–13 (2018).

43. Varoqueaux, F. et al. Total arrest of spontaneous and evoked synaptic transmission but normal synaptogenesis in the absence of Munc13-mediated vesicle priming. Proc Natl Acad Sci U S A 99, 9037–9042 (2002).

44. Richmond, J. E., Davis, W. S. & Jorgensen, E. M. UNC-13 is required for synaptic vesicle fusion in C. elegans. Nat Neurosci 2, 959–964 (1999).

45. Böhme, M. A. et al. Rapid active zone remodeling consolidates presynaptic potentiation. Nat Commun 10, (2019).

46. Böhme, M. A. et al. Active zone scaffolds differentially accumulate Unc13 isoforms to tune Ca(2+) channel-vesicle coupling. Nat Neurosci 19, 1311–1320 (2016).

47. Blaum, N. et al. Monoamine-induced diacylglycerol signaling rapidly accumulates Unc13 in nanoclusters for fast presynaptic potentiation. bioRxiv 2025.01.10.632340 (2025) doi:10.1101/2025.01.10.632340.

48. Fulterer, A. et al. Active Zone Scaffold Protein Ratios Tune Functional Diversity across Brain Synapses. Cell Rep 23, 1259–1274 (2018).

49. Dittman, J. S. Unc13: a multifunctional synaptic marvel. Curr Opin Neurobiol 57, 17– 25 (2019).

50. Michelassi, F., Liu, H., Hu, Z. & Dittman, J. S. A C1-C2 Module in Munc13 Inhibits Calcium-Dependent Neurotransmitter Release. Neuron 95, 577–590.e5 (2017).

51. Taschenberger, H., Woehler, A. & Neher, E. Superpriming of synaptic vesicles as a common basis for intersynapse variability and modulation of synaptic strength. Proc Natl Acad Sci U S A 113, E4548–E4557 (2016).

52. Rhee, J. S. et al. β phorbol ester- and diacylglycerol-induced augmentation of transmitter release is mediated by Munc13s and not by PKCs. Cell 108, 121–133 (2002).

53. Betz, A. et al. Munc13-1 is a presynaptic phorbol ester receptor that enhances neurotransmitter release. Neuron 21, 123–136 (1998).

54. Basu, J., Betz, A., Brose, N. & Rosenmund, C. Munc13-1 C1 domain activation lowers the energy barrier for synaptic vesicle fusion. J Neurosci 27, 1200–1210 (2007).

55. Root, C. M. et al. A presynaptic gain control mechanism fine-tunes olfactory behavior. Neuron 59, 311–321 (2008).

56. Qiu, Y. et al. Natural environment statistics in the upper and lower visual field are reflected in mouse retinal specializations. Current Biology 31, 3233–3247.e6 (2021).

57. Ramirez, L. & Dickman, R. Data-Driven Models of Efficient Chromatic Coding in the Outer Retina. eNeuro 9, (2022).

58. Olshausen, B. A. & Field, D. J. Emergence of simple-cell receptive field properties by learning a sparse code for natural images. Nature 381, 607–609 (1996).

59. Ganguli, D. & Simoncelli, E. P. Efficient sensory encoding and Bayesian inference with heterogeneous neural populations. Neural Comput 26, 2103–2134 (2014).

60. Keller, A. & Vosshall, L. B. Influence of odorant receptor repertoire on odor perception in humans and fruit flies. Proc Natl Acad Sci U S A 104, 5614–5619 (2007).

61. Zwicker, D., Murugan, A. & Brenner, M. P. Receptor arrays optimized for natural odor statistics. Proc Natl Acad Sci U S A 113, 5570–5575 (2016).

62. Tesileanu, T., Cocco, S., Monasson, R. & Balasubramanian, V. Adaptation of olfactory receptor abundances for efficient coding. Elife 8, (2019).

63. Cognigni, P., Felsenberg, J. & Waddell, S. Do the right thing: neural network mechanisms of memory formation, expression and update in Drosophila. Curr Opin Neurobiol 49, 51–58 (2017).

64. Ling, D., Zhang, L., Saha, D., Chen, A. B. & Raman, B. Adaptation invariant concentration discrimination in an insect olfactory system. Elife 12, (2023).

65. Gomez-Marin, A., Stephens, G. J. & Louis, M. Active sampling and decision making in Drosophila chemotaxis. Nature Communications 2011 2:1 2, 1–10 (2011).

66. Gepner, R., Skanata, M. M., Bernat, N. M., Kaplow, M. & Gershow, M. Computations underlying Drosophila phototaxis, odor-taxis, and multi-sensory integration. Elife 4, (2015).

67. Vickers, N. J. Mechanisms of animal navigation in odor plumes. 10.2307/1542524 198, 203–212 (2000).

68. Murlis, J., Elkinton, J. S. & Cardé, R. T. Odor plumes and how insects use them. Annu Rev Entomol 37, 505–532 (1992).

69. Kadakia, N. et al. Odour motion sensing enhances navigation of complex plumes. Nature 2022 611:7937 611, 754–761 (2022).

70. Dolan, M. J. et al. Neurogenetic dissection of the drosophila lateral horn reveals major outputs, diverse behavioural functions, and interactions with the mushroom body. Elife 8, (2019).

71. Frechter, S. et al. Functional and anatomical specificity in a higher olfactory centre. Elife (2019) doi:10.7554/eLife.44590.

72. Taisz, I. et al. Generating parallel representations of position and identity in the olfactory system. Cell 186, 2556–2573.e22 (2023).

73. Kim, H. S., Santana, G. M., Sancer, G., Emonet, T. & Jeanne, J. M. Divergent synaptic dynamics originate parallel pathways for computation and behavior in an olfactory circuit. Current Biology 35, 3146–3162.e8 (2025).

74. Álvarez-Salvado, E. et al. Elementary sensory-motor transformations underlying olfactory navigation in walking fruit-flies. Elife 7, e37815 (2018).

75. Murmu, M. S., Stinnakre, J. & Martin, J. R. Presynaptic Ca2+ stores contribute to odor-induced responses in Drosophila olfactory receptor neurons. Journal of Experimental Biology (2010) doi:10.1242/jeb.046474.

76. Murmu, M. S., Stinnakre, J., Réal, E. & Martin, J. R. Calcium-stores mediate adaptation in axon terminals of Olfactory Receptor Neurons in Drosophila. BMC Neurosci (2011) doi:10.1186/1471-2202-12-105.

77. Silva, M., Tran, V. & Marty, A. Calcium-dependent docking of synaptic vesicles. Trends Neurosci 44, 579–592 (2021).

78. Rozenfeld, E., Ehmann, N., Manoim, J. E., Kittel, R. J. & Parnas, M. Homeostatic synaptic plasticity rescues neural coding reliability. Nature Communications 2023 14:1 14, 1–14 (2023).

79. Ramirez, L. Trade-off between coding efficiency and color space in outer retinal circuits with colored oil droplets. Vision Res 208, (2023).

80. Baden, T. The vertebrate retina: a window into the evolution of computation in the brain. Curr Opin Behav Sci 57, (2024).

81. Wang, X., Roberts, P. A., Yoshimatsu, T., Lagnado, L. & Baden, T. Amacrine cells differentially balance zebrafish color circuits in the central and peripheral retina. Cell Rep 42, 112055 (2023).

82. Brand, P. et al. The origin of the odorant receptor gene family in insects. Elife 7, (2018).

83. Ruta, V. et al. A dimorphic pheromone circuit in Drosophila from sensory input to descending output. Nature 468, 686–690 (2010).

84. Stensmyr, M. C. et al. A conserved dedicated olfactory circuit for detecting harmful microbes in drosophila. Cell 151, 1345–1357 (2012).

85. Kondo, S. & Ueda, R. Highly improved gene targeting by germline-specific Cas9 expression in Drosophila. Genetics 195, 715–721 (2013).

86. Nagel, K. I., Hong, E. J. & Wilson, R. I. Synaptic and circuit mechanisms promoting broadband transmission of olfactory stimulus dynamics. Nature Publishing Group 18, 56–65 (2015).

87. Chou, Y. H. et al. Diversity and wiring variability of olfactory local interneurons in the Drosophila antennal lobe. Nat Neurosci (2010) doi:10.1038/nn.2489.

