## Supplementary material for "Undoing of firing rate adaptation enables invariant population codes"

**Table S1: Key resources**

| **Reagent or Resource** | **Source** | **Identifier** |
| --- | --- | --- |
| Chemicals | | |
| Methyl acetate | Sigma-Aldrich Co | Art. #45999, CAS 79-20-9 |
| 2-butanone | Carl Roth | Art. #T920.1, CAS 78-93-3 |
| Mineral Oil | Sigma-Aldrich Co | Art. #330779, CAS 8042-47-5 |
| CGP 54626 hydrochloride | Tocris Bioscience | Art. #1088, CAS 149184-21-4 |
| Picrotoxin | Sigma-Aldrich Co | Art. #P1675, CAS 124-87-8 |
| Atropine | Sigma-Aldrich Co | Art. #A0132, CAS 51-55-8 |
| Tubocurarine | Sigma-Aldrich Co | Art. #T2379, CAS 6989-98-6 |
| Imidacloprid | Sigma-Aldrich Co | Art. #37894, CAS 138261-41-3 |
| All trans retinal | Sigma-Aldrich Co | Art. #R2500, CAS 116-31-4 |
| Bondic cartridge with Bondic liquid plastic | Bondic | Art. #BONKART |
| Bondic UV-LED | Bondic | Art. #BONLED |
| Experimental Models: Organisms/Strains | | |
| D. melanogaster: or42b-Gal4 | Bloomington Drosophila Stock Center (BDSC) | RRID: BDSC_9972 |
| D. melanogaster: or10a-Gal4 | BDSC | RRID: BSDC_9944 |
| D. melanogaster: GH146-Gal4 | Gift from A. Fiala | Gift from A. Fiala |
| D. melanogaster: orco-Gal4 | BDSC | RRID: BSDC_23292 |
| D. melanogaster: UAS-GCaMP6f | BDSC | RRID: BSDC_42747 |
| D. melanogaster: UAS-CsChrimson | BDSC | RRID: BSDC_82181 |
| D. melanogaster: UAS- shibire$^{ts}$ | Gift from M. Silies | |
| D. melanogaster: UAS-ASAP2s | Gift from T. Clandinin | |
| D. melanogaster: lexAop-GCaMP6f | BDSC | RRID: BSDC_44277 |
| D. melanogaster: orco-LexA | Gift from H. Scholz | |
| Software and algorithms | | |
| Fiji | | RRID: SCR_002285 |
| MATLAB | MathWorks | RRID: SCR_001622 |
| Adobe Illustrator | Adobe | RRID: SCR_010279 |

#### Table S2: Genetic *D. melanogaster* strains

| Name | Genotype | Figure |
|---|---|---|
| *GH146 > GCaMP6f* | *w-; GH146-Gal4/UAS-GCaMP6f ; +* | 6a-c, 6e, S4a-c |
| *GH146 > ASAP2s* | *w+/w-; GH146-Gal4/UAS-ASAP2s ; +* | 6d, S4d,e |
| *or42b > GCaMP6f, csChrimson* | *w+/w-; UAS-GCaMP6f/UAS-CsChrimson; or42b-Gal4/+* | 3b-d |
| *or42b > GCaMP6f, shibire*$^{ts}$ | *w+/w-; UAS-GCaMP6f/+; or42b-Gal4/UAS-shibire*$^{ts}$ | 4a-f |
| *or42b > GCaMP6f (control)* | *w+/w-; UAS-GCaMP6f/+; or42b-Gal4/+* | 4a-f |
| *or10a > GCaMP6f, shibire*$^{ts}$ | *w-; or10a-Gal4/UAS-GCaMP6f; UAS-shibire*$^{ts}$*/+* | S1a-d |
| *or10a > GCaMP6f (control)* | *w-; or10a-Gal4/UAS-GCaMP6f; +* | S1a-d |
| *GH146 >shibire*$^{ts}$*;* *orco > GCaMP6f* | *w-; GH146-Gal4/lexAop-GCaMP6f ; orco-LexA/UAS-shi[ts]* | S1e-f |
| *GH146; orco > GCaMP6f (control)* | *w-; GH146-Gal/lexAop-GCaMP6f ; orco-LexA/TM2* | S1e-f |
| *orco > GCaMP6f* | *w+/w-; UAS-GCaMP6f/+; orco-GAL4/+* | 4g-n, 7b, S2 |
| *GH146 > GCaMP6f* | *w-; GH146-Gal4/UAS-GCaMP6f ; +* | 7c |
| *GH146 > GCaMP6f; Unc13H1723K* | *w-; GH146-Gal4/UAS-GCaMP6f ; +; Unc13H1723K* | 7d |

#### Table S3: Airflow dilutions for odor

| C= test stimulus B = background | Odor airflow (sccm) | Equivalent dilution |
|---|---|---|
| C4/B4 | 0.03 | $10^{-4.5}$ |
| C5/B5 | 0.10 | $10^{-4}$ |
| C6/B6 | 0.30 | $10^{-3.5}$ |
| C7/B7 | 1 | $10^{-3}$ |
| C8/B8 | 3 | $10^{-2.5}$ |
| C9/B9 | 10 | $10^{-2}$ |

### Supplementary Figures

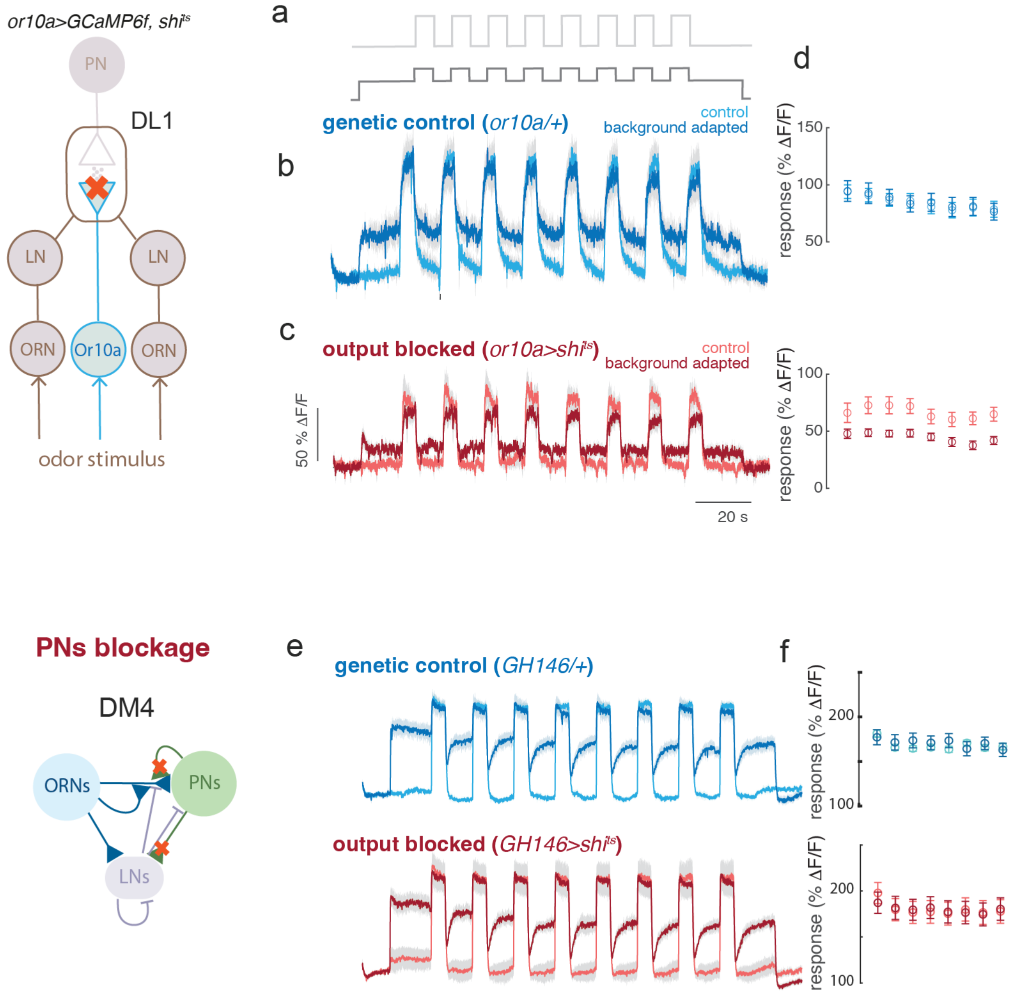


**Figure S1: Role of synaptic release for ORN background invariance.**
**a)** Genetic blockage of Or10a-ORNs neurotransmitter release. Stimulation protocol in control (light gray) and background adapted conditions (dark gray). **b)** Calcium response of the DL1 glomerulus in genetic control flies showing background-invariant responses. **c)** Calcium response in flies with blocked ORN synapses showing background dependent responses. **d)** Quantification of peak response for each of the 8 test pulses. Error bars indicate sem (N=4, 7).
**e)** Blocking of PN synaptic output does not affect ORN presynaptic calcium responses. Traces represent mean calcium responses of DM4 ORNs in control (light colors) and background adapted conditions (dark colors) for control (blue) and test flies (red). Test flies express shi$^{ts}$ under control of GH146-GAL4 that labels PNs. **f)** Quantification of peak response for each of the 8 test pulses. Error bars indicate sem.

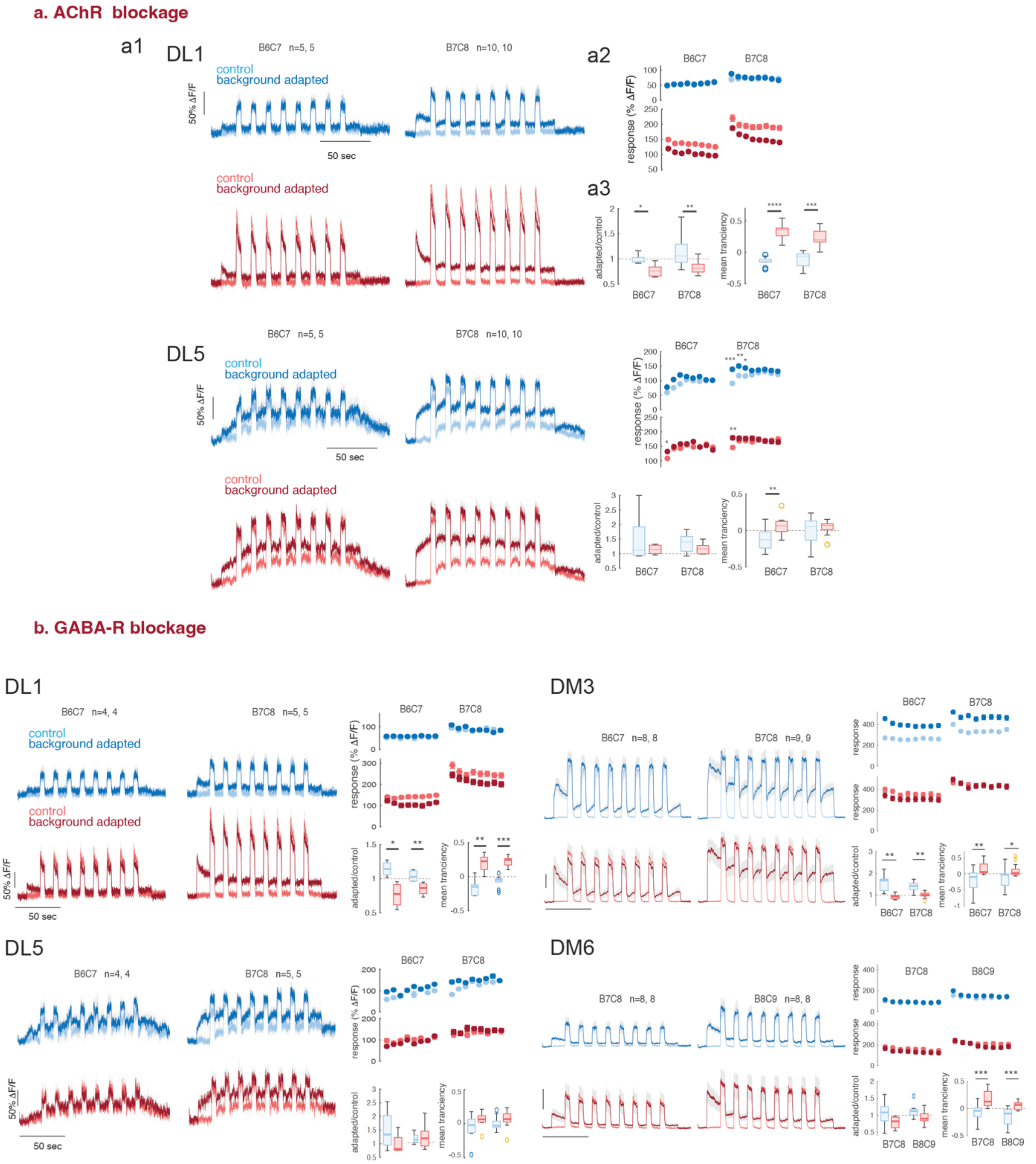


**Figure S2: Glomerulus specific mechanisms for presynaptic calcium modulation**.

**a)** Pharmacological blockage of AchRs (100 µM atropine + 20 µM tubocurarine + 200 nM imi, see Methods). **a1)** Traces represents mean calcium responses of DL1 ORNs in control (light colors) and background adapted conditions (dark colors) before (blue) and after (red) drug application. **a2)** Quantification of peak response for each of the 8 test pulses. Error bars indicate SEM but might not be visible. **a3)** Boxplot quantification of the degree of adaptation and response transiency, indicating median, quartiles max/min and outliers. Same organization for glomerulus DL5.

**b)** Pharmacological blockage of GABA receptors (25 µM CGP54626 + 5 µM picrotoxin, see Methods) in 4 glomeruli. Same organization of the plots as in **a)**. 2-butanone was used for DL1 and DL5 and butyl acetate for DM3 ($10^{-2}$ liquid dilution in PO) and DM6 ($10^{-3}$ liquid dilution in PO). *p<0.05, **p<0.01, ***p<0.001. Sample size is indicated for each dataset in the figure.

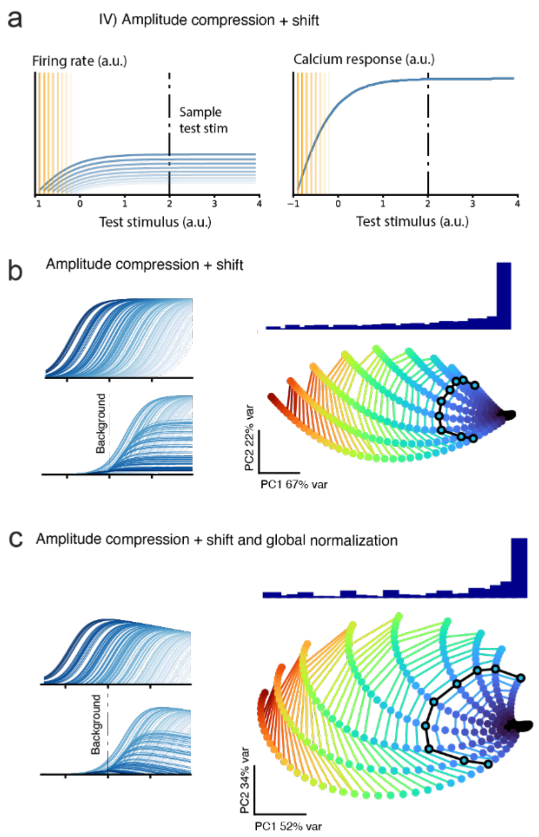


**Figure S3: Population of ORNs with divisive normalization. a)** Firing rate and calcium response for an adaptive function combining amplitude-compression and amplitude-shift as predicted by a model of background compensation and response normalization (Appendix 1). As in Fig. 5, Orange lines indicate the value of the adaptive background stimulus. Curves with different shades of red/blue correspond to different adapted conditions. Dotted lines indicate a representative test stimulus. **b)** 2D-manifold of the ORN population response to stimuli of different ON (color coded) and OFF (black) contrast in different background adapted conditions. The histogram represents the distribution of population states calculated on the first principal component. Same as plots b) and e) in Fig. 8 but now for the combined amplitude adaptation mechanism.

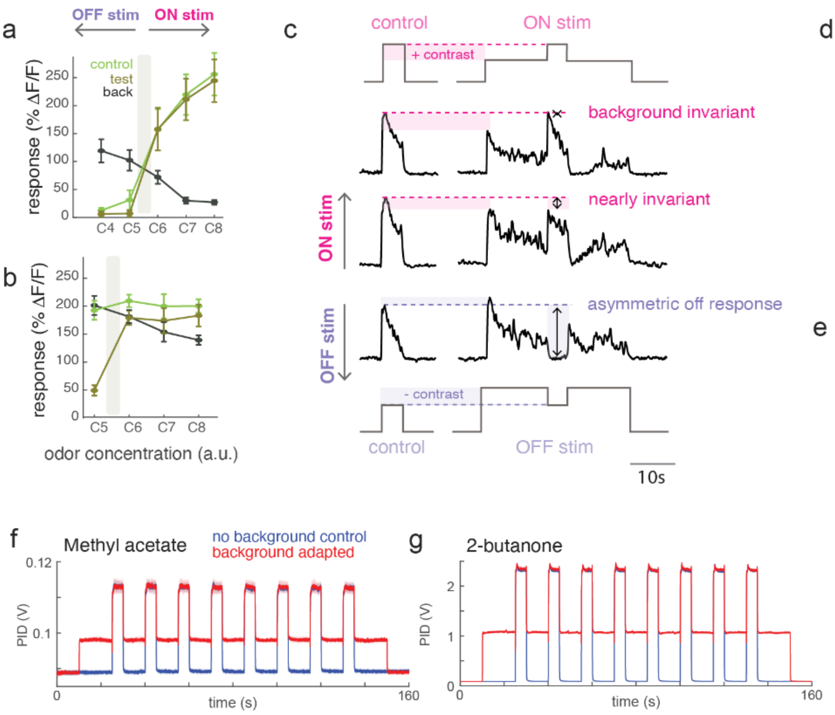


**Figure S4: PNs response to ON and OFF stimuli**. To cover a range of ON and OFF responses of PNs, we conducted two sets of experiments using different liquid solutions of the odors. To generate main **Fig. 6c** we analyzed the single trials of these two datasets. Here we show mean responses (**a, b**) and some examples of response to small contrast stimuli (**c**).

**a-b)** Peak calcium response to different combinations of ON, OFF and background stimuli using methyl acetate. *Green*: mean response to non-adapted control. *Black*: mean response to background stimulus. *Dark green*: mean response to the same stimulus as in green after 15 seconds of background adaptation. Error bars indicate SEM, the shaded area indicated the stimulus that equals the background concentration. Liquid dilution of methyl acetate for the background was $10^{-4}$ in **a)** and $10^{-3}$ in **b)**, therefore the higher background response in **b)**. The liquid dilution for the pulse was $10^{-2}$ in **a)** and $10^{-3}$ in **b)**. Labels on the x-axis corresponds to gas phase dilutions generated by the odor delivery system as indicated in **table S3**.

**c)** Three single trials from the dataset showing responses to ON and OFF stimuli with small contrast. The shaded area indicates the difference in response between test and background stimulus. The dotted line indicates the response to the control stimulus. These traces show how a small positive contrast leads to nearly perfectly invariant response (same as controls), while a small negative contrast leads to a strong decrease in calcium, much lower than the test stimulus amplitude. **d)** Control and background adapted response measure in DM2 using either a calcium or voltage sensor. Thick lines indicate the mean ΔF/F and the shaded areas sem. Liquid dilution of methyl acetate is $10^{-4}$ for the background and $10^{-2}$ for the pulse. **e)** Boxplot representing median, quartiles, min/max values and outliers of ASAP2s signals from two glomeruli. (DM2 N=19, VM7d N=23, *$p<0.05$, Kruskal-Wallis).

**f)** Average PID response to methyl acetate using a liquid dilution of $10^{-2}$ and background airflow dilution of C6 and pulse C7 (N=4) with stimulation protocol in control (blue) and background adapted conditions (red). **g)** Same as d) for pure 2-butanone with a background airflow dilution of C7 and pulse C8 (N=4).

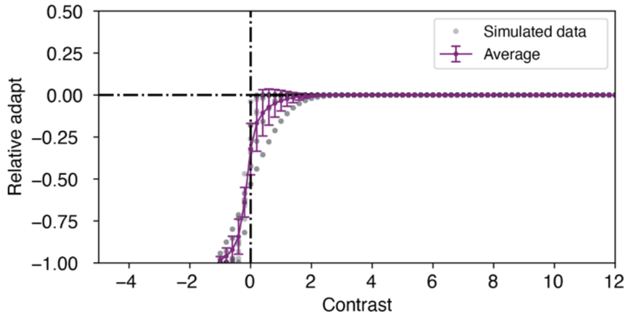


**Figure S5: Modeling PNs relative adaptation for different contrast stimuli.** We model the responses of PN neurons in a background adapted condition as a noisy process with probability $\mathrm{N}(s - s_B, \sigma)$, where $s_B$ is the background stimulus and $\sigma$ is the noise std. We simulate PN responses across different contrast stimuli and show that a standard deviation of $\sigma = 0.5$ fits the experimental observations shown in **Fig. 6c**. For each contrast value, we average the relative adaptation over a set of 10 different background conditions ranging from -4 to 4 in the stimulus log. scale.

#### Appendix 1

#### Adaptation model: firing rate and calcium response with divisive normalization

In the main text we have modeled the dynamics of calcium responses in ORNs as a linear function of their firing rate and the inhibitory inputs from LNs (**Fig. 5** and Methods). However, previous work by Olsen et al. [1] has shown that LN inhibition in the AL performs a divisive (rather than linear) transformation from ORNs to PNs firing rates. Therefore, we ask if a divisive model would also be able to support the background compensation. It should be noted that the divisive model was not originally proposed to fit the transformation between ORN firing rate and ORN axonal calcium. Moreover, the model is based on mean responses and does not capture response dynamics nor considers adaptive processes. Nonetheless, in the following we show that also when assuming a divisive effect of LN activity on ORN calcium, background invariance can only be achieved if the firing rate adapts in the amplitude domain, rather than by a shift in sensitivity.

We consider the calcium response of ORNs without odor background to be:

$$R[F(s)] = \frac{F(s)}{F(s) + \sigma} \qquad (s.1)$$

This relationship implies that the effect of inhibition is not linear (as in Eq. 3 of the main text) but dependents on the input stimulus: $\Delta_I = 1/(F(s) + \sigma)$ with $\sigma$ a constant. Background invariance is achieved if $R[F(s, s_B)] = R[F(s)] = R[F(s_B)] + R[F'(s)]$ (Eq. 4 of main text). Using Eq. (s.1), we obtain:

$$\frac{F'(s)}{L(F'(s))} = \frac{F(s)}{F(s) + \sigma} - \frac{F(s_B)}{F(s_B) + \sigma} \qquad (s.2)$$

where we assume a general form of the normalization factor $\mathrm{L}(F'(s))$ in adapted conditions. For equation s.2 to hold, it is required both that:

$$F'(s) = \sigma \cdot (F(s) - F(s_B))$$

and, using s.3 in s.2, that:

$$L(F'(s)) = A \cdot F'(s) + \sigma'$$

with $A = \frac{F(S_B)+\sigma}{\sigma}$ and $\sigma' = (F(s_B) + \sigma)^2$. Eq. (s.3) indicates that to achieve background invariance the adapted firing rate should be a combination of an amplitude compression (first term) and an amplitude shift (second term), as we obtained in the case of the dynamic linear model (Eq. 7, 8 in main text). In addition, Eq. (s.4) indicates that background invariance with divisive normalization requires adaptation of the LN feedback and therefore a background dependent normalization term. Specifically, in adapted conditions the normalization factor is still linear but a stronger effect ($A > 1$ $and$ $\sigma' > \sigma$)

Including Eq. (1) into Eq. (s.4), we find the update rules,

$$\begin{aligned} \beta' &= \beta \\ \alpha' &= \alpha\big[1 + R_{Ca}[F(s_B)]\big] \\ \gamma' &= \gamma + \frac{\sigma R_{Ca}[F(s_B)]}{\alpha(1 + R_{Ca}[F(s_B)])} \end{aligned} \qquad (s.5)$$

This shows that the combined adaptation mechanism depends uniquely on the background adaptation response in both ORNs and LNs. Furthermore, it satisfies the calcium invariant condition. As done for the main dynamic model, we studied the consequences of this combined adaptation mechanism for single ORNs and for the population responses. We show that background invariance is indeed preserved for all test odors similarly to the ORNs implementing an amplitude-based adaptation (**Fig. S3a**). Regarding the encoding of ORNs at the population level, we performed the same analysis shown in Fig. 8, and show that the combination of both amplitude -shift and -compression mechanisms also preserve information about input contrast and lead to ON-background invariant responses in populations of PN neurons (**Fig. S3b**).

1. Olsen, S. R., Bhandawat, V. & Wilson, R. I. Divisive normalization in olfactory population codes. *Neuron* 66, 287–299 (2010).
